# Determinants of raffinose family oligosaccharide use in *Bacteroides* species

**DOI:** 10.1101/2024.06.07.597959

**Authors:** Anubhav Basu, Amanda N.D. Adams, Patrick H. Degnan, Carin K. Vanderpool

## Abstract

*Bacteroides* species are successful colonizers of the human gut and can utilize a wide variety of complex polysaccharides and oligosaccharides that are indigestible by the host. To do this, they use enzymes encoded in Polysaccharide Utilization Loci (PULs). While recent work has uncovered the PULs required for use of some polysaccharides, how *Bacteroides* utilize smaller oligosaccharides is less well studied. Raffinose family oligosaccharides (RFOs) are abundant in plants, especially legumes, and consist of variable units of galactose linked by ⍺-1,6 bonds to a sucrose (glucose ⍺-1-β-2 fructose) moiety. Previous work showed that an α-galactosidase, BT1871, is required for RFO utilization in *Bacteroides thetaiotaomicron*. Here, we identify two different types of mutations that increase *BT1871* mRNA levels and improve *B. thetaiotaomicron* growth on RFOs. First, a novel spontaneous duplication of *BT1872* and *BT1871* places these genes under control of a ribosomal promoter, driving high *BT1871* transcription. Second, nonsense mutations in a gene encoding the PUL24 anti-sigma factor likewise increase *BT1871* transcription. We then show that hydrolases from PUL22 work together with BT1871 to break down the sucrose moiety of RFOs and determine that the master regulator of carbohydrate utilization (BT4338) plays a role in RFO utilization in *B. thetaiotaomicron*. Examining the genomes of other *Bacteroides* species, we found homologs of BT1871 in subset and show that representative strains of species containing a BT1871 homolog grew better on melibiose than species that lack a BT1871 homolog. Altogether, our findings shed light on how an important gut commensal utilizes an abundant dietary oligosaccharide.

**Importance:** The gut microbiome is important in health and disease. The diverse and densely populated environment of the gut makes competition for resources fierce. Hence, it is important to study the strategies employed by microbes for resource usage. Raffinose family oligosaccharides are abundant in plants and are a major source of nutrition for the gut microbiota since they remain undigested by the host. Here, we study how the model gut commensal, *Bacteroides thetaiotaomicron* utilizes raffinose family oligosaccharides. This work highlights how an important member of the microbiota uses an abundant dietary resource.

## Introduction

The gut microbiome plays an important role in human health and development by facilitating energy extraction from food (1), synthesis of vitamins (2, 3), protection from pathogen colonization (4, 5), enhancing the immune system (6, 7), and modulating gut-brain communication (8). Although diverse, the human gut microbiome is dominated by two major phyla, namely the *Bacillota* and *Bacteroidota* (9, 10). Members of the *Bacteroidota* owe their success to their ability to break down and use complex host and plant-derived polysaccharides in the diet. They accomplish this using paralogous gene clusters known as Polysaccharide Utilization Loci (PUL) that encode glycan sensing, transport, and degrading enzymes (11–15). The prototypical PUL is characterized by the presence of homologs of SusC (an outer membrane oligosaccharide transporter) and SusD (an outer membrane oligosaccharide binding protein) found in the Starch Utilization System (Sus) (16–18). They also contain other membrane bound and periplasmic proteins required to bind, break down and transport complex carbohydrates (19, 20).

Each PUL is specialized for degrading distinct polysaccharides and hence PUL gene expression is tightly regulated. Hybrid Two Component Systems (HTCS) and extracytoplasmic function (ECF) sigma factor/anti-sigma pairs are the most common regulators of PULs in *Bacteroides* (21, 22). HTCS combine the sensor kinase and response regulator proteins of a classical two component system into a single polypeptide spanning the inner membrane. The recognition domain of the HTCS senses a unique oligosaccharide signal usually 2-8 subunits in length which causes sensor domain autophosphorylation and transfer of the phosphate to the regulator domain that promotes transcription of target genes (21, 23–28). PULs regulated by ECF sigma/anti-sigma pairs are usually associated with breakdown of host derived polysaccharides (29). Transport of oligosaccharides through the SusC-like protein is coupled to conformational changes in the anti-sigma, resulting in release of the cognate sigma factor from the inner membrane and transcription of target genes (14, 30).

*Bacteroides thetaiotaomicron*, a model gut microbe, has over a 100 PULs and dedicates ∼18% of its genome to encoding functions for carbohydrate sensing and usage (31, 32). In recent years, many studies have probed the genetic and mechanistic details of how long chain polysaccharides are broken down and utilized *B. thetaiotaomicron* (27–29, 33–36). However, there is less work focused on how *B. thetaiotaomicron* or other *Bacteroides* species can use smaller oligosaccharides. One such group of oligosaccharides is the raffinose family oligosaccharides (RFOs). RFOs are soluble carbohydrates and are functionally ⍺-D-galactosyl derivatives of sucrose, a disaccharide of glucose and fructose (Fig. S1 and (37)). Raffinose is the simplest RFO and is a trisaccharide where a single galactose is ⍺-1,6 linked to the glucose moiety of sucrose. Longer RFOs such as stachyose and verbascose contain two and three galactose residues, respectively (Fig. S1). RFOs are highly abundant in the seeds of many crops, particularly in the members of the legume family such as soybean, lentils, and chickpea (37–39). In plants, RFOs function in storage and translocation (40–42), stress tolerance (43–45), seed germination desiccation tolerance (46–49). RFOs are indigestible by humans since we lack the ⍺-galactosidases required to break them down (50). Hence, RFOs in the diet pass undigested to the colon where they are utilized by a variety of microbes (51–53).

Recent studies have revealed the importance of RFOs in human health as prebiotics because they can modulate the abundance of beneficial bacteria in the gut (51, 52, 54).

Because of their ability to influence the microbiota, it is important to elucidate the mechanism by which gut bacteria can utilize RFOs. Various *Bifidobacterium* species, *B. subtilis* and *S. pneumoniae* contain dedicated operons for sensing, transporting, and breaking down RFOs (53, 55–57). Other microbes such as *E. coli and E. faecium* harbor these functionalities on plasmids (58, 59). However, homologues of these systems are absent in *Bacteroides* species. Previous work from our lab found that an ⍺-galactosidase encoded by *BT1871* in PUL24 of *B. thetaiotaomicron* is important for RFO utilization *in vitro* (60). Deletion of *BT1871* led to decreased growth on RFOs as the sole carbon source.

In this study, we show that the efficiency of RFO use in *B. thetaiotaomicron* is limited by low levels of expression of *BT1871*, encoding an ⍺-galactosidase that breaks the α-1,6 bond between glucose and galactose in RFOs. We found two different types of mutations that promote higher *BT1871* transcription to increase growth on RFOs. First, we serendipitously identified *B. thetaiotaomicron* strains with a novel duplication involving *BT1871* that leads to substantially better growth on RFOs compared to strains with only a single copy of *BT1871*.

Second, we demonstrated that disruption of *BT1876,* encoding the PUL24 anti-sigma factor also increases *BT1871* transcript levels and leads to better RFO utilization. We then established that full RFO degradation by *B. thetaiotaomicron* requires the PUL24 α-galactosidase BT1871 as well as sucrases encoded by genes in PUL22. Investigation of regulatory mechanisms controlling RFO use revealed that the master regulator of carbohydrate utilization, BT4338, is required for growth on RFOs through control of the expression of *BT1871*. Finally, we show that *BT1871* homologs in other *Bacteroides* species are also important for their ability to use the disaccharide melibiose. Taken together, our findings reveal key players that help *B. thetaiotaomicron* and other *Bacteroides* species utilize RFOs.

## Results

### A novel duplication of *BT1871* confers a growth advantage to *B. thetaiotaomicron* growing on RFOs

Previously, we found that loss of the RNA binding protein RbpB (encoded by *BT1887*) in *B. thetaiotaomicron* caused a growth defect on raffinose family oligosaccharides (RFOs) as the sole carbon source (60). This phenotype was linked to the reduced expression of the *BT1871- BT1872* locus in the *rbpB* mutant strain compared to the parent strain. Importantly, *BT1871* is an α-galactosidase capable of breaking down RFOs and its deletion also caused growth defects on RFOs (60).

During construction of *rbpB* mutations in different backgrounds, we noticed that some mutant isolates had no growth defects on RFOs while other mutants showed growth defects similar to the original *rbpB* mutant. To better understand this phenotypic variability, we performed long- and short-read whole genome sequencing of the two types of mutant isolates and the parent strain. We found that the parent strain had a duplication of the *BT1871-BT1872* locus which is not present in the NCBI reference genome (NC_004663.1). This duplication involves the promoter and 5’ end of a 16S rRNA gene, an IS3 transposable element (*BT1869- BT1870*) and *BT1871* and *BT1872* (Fig. 1A). The duplication places *BT1871* and *BT1872* downstream of a strong ribosomal promoter. Once we had defined the structure of the duplication, we were able to isolate wild-type strain derivatives that had lost the duplication. Comparing transcript levels of *BT1871* in strains with (dupl+) or without (<u>dupl-)</u> the duplication we found that dupl+ strains had substantially higher *BT1871* transcript levels (Fig. 1B, WT dupl+ and Δ*rbpB* dupl+) compared to the strains without the duplication (Fig. 1B, WT dupl- and Δ*rbpB* dupl-). Importantly, the phenotypes we observed previously, including growth on the RFO subunit melibiose, that we previously attributed to loss of *rbpB,* were instead caused by the loss of the duplication. This is evident when comparing the growth of the wild-type (dupl+) and two *rbpB* mutant isolates, one with (dupl+) and one without (dupl-) the duplication, on melibiose (Fig. 1C). Both wild-type dupl+ and *rbpB* dupl+ strains grew well on melibiose (Fig. 1C), consistent with their high levels of expression of *BT1871* (Fig. 1B), which encodes the α-galactosidase that breaks the melibiose disaccharide bond (61, 62). In contrast, the wild-type dupl- and *rbpB* dupl- strains grew more slowly on melibiose (Fig. 1C) consistent with greatly reduced levels of *BT1871* expression in these strains (Fig. 1B). Hence, we conclude that the growth phenotype of *B. thetaiotaomicron* on RFOs is linked to the duplication status and expression of the *BT1871* gene and is not linked to the function of RbpB.

**Figure 1:**
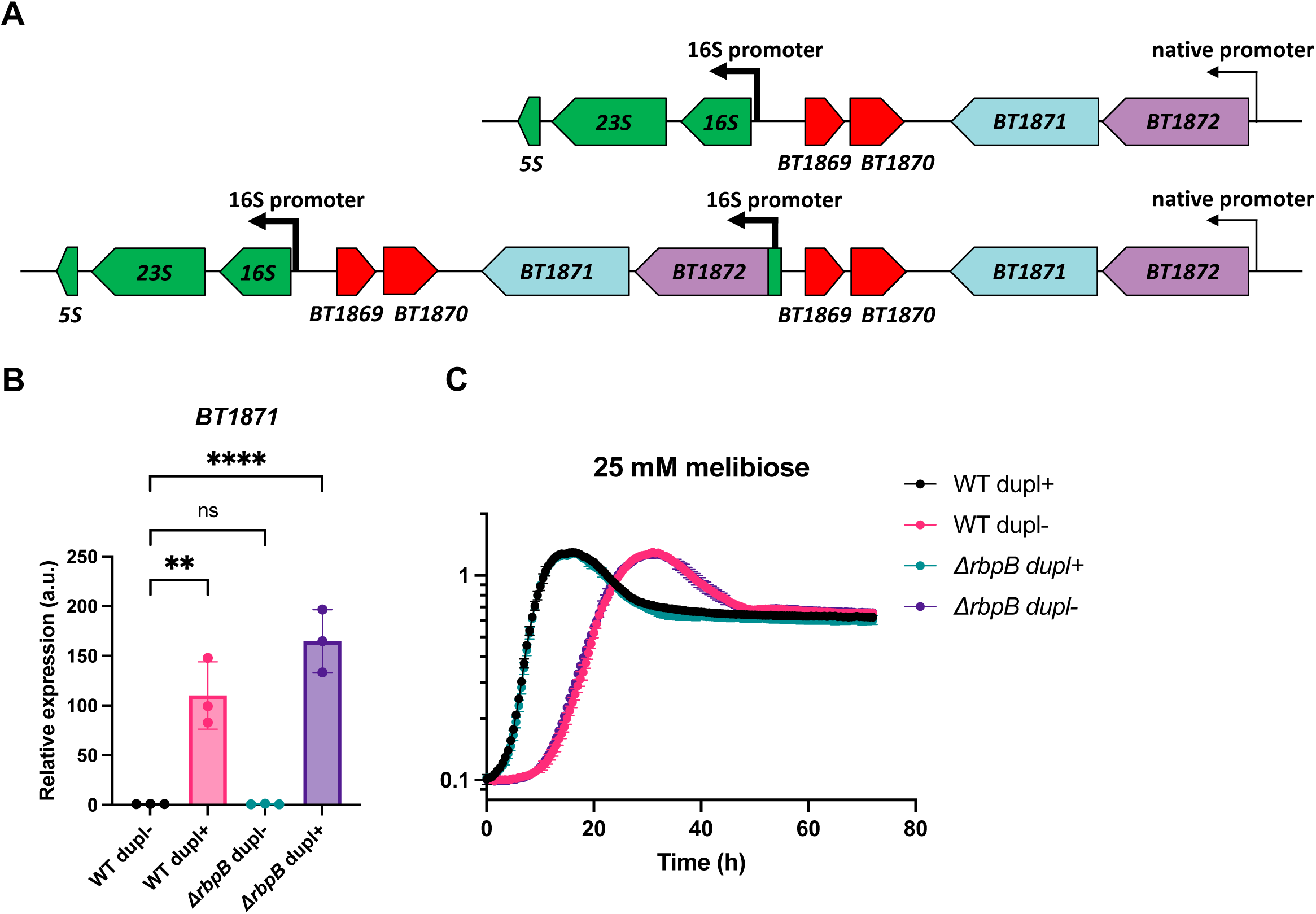
Duplication of the *BT1871-BT1872* locus provides *B. thetaiotaomicron* a growth advantage on melibiose. (A) Structure of the *BT1871-BT1872* duplication in a dupl+ strain (bottom) compared to a strain without the duplication (dupl-, top). Red gene models represent genes in an IS3 element while green gene models represent genes in a ribosomal RNA operon. (B) qRT-PCR to measure of *BT1871* mRNA levels in the indicated strains grown to mid-log phase in rich (TYG) media. The bars represent the mean and SD of n=3 biological replicates. All values are normalized to the level in WT dupl-. Statistical significance was determined by one way ANOVA. **, P < 0.01; ****, P < 0.0001. (C) Growth curves of the indicated strains on minimal media with melibiose as the sole carbon source. The points and error bars represent the mean and SD of n=3 biological replicates.

### Mutations in the anti-sigma gene of PUL24 confer better RFO utilization in *B. thetaiotaomicron* through increased transcription of *BT1871*

To understand more about the role of *BT1871* in carbon source use and determine whether the duplication impacts use of carbon sources beyond RFOs, we monitored the growth of wild-type dupl+ and dupl- cells on a variety of carbon sources. We identified the sugar ⍺-methyl galactoside (AMG) as a carbon source that supported slow growth of dupl+ cells while dupl- cells could not grow even after 72 hours of incubation (Fig. 2A). Since AMG has the same galactose-⍺-1,6 bond found in melibiose and RFOs and *BT1871* is important for growth on RFOs, we hypothesized that the higher levels of BT1871 in the dupl+ strain were responsible for the observed phenotype on AMG.

**Figure 2.**
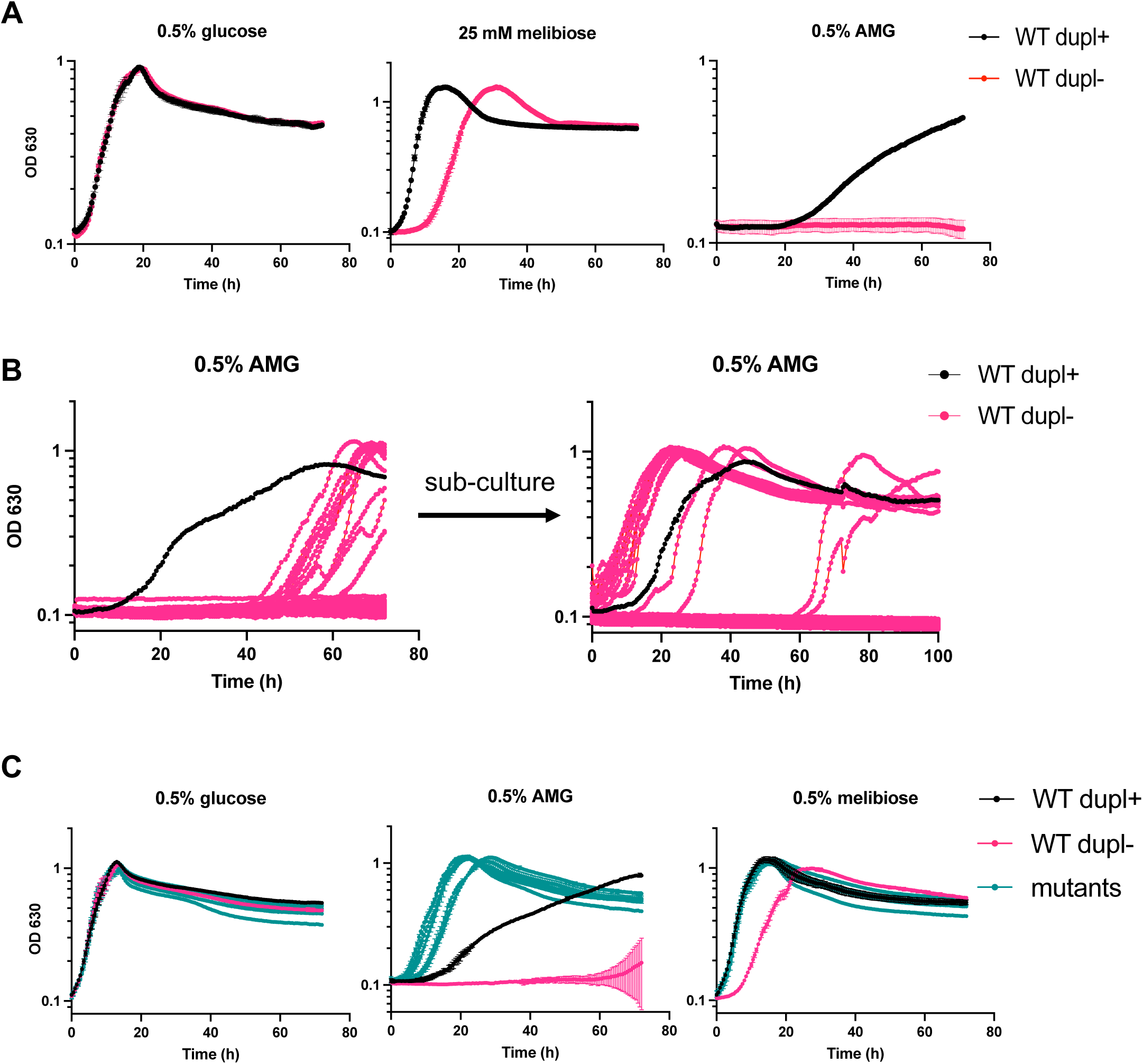
*B. thetaiotaomicron* mutants derived from WT dupl- cells grow on AMG. (A) Growth curves of WT dupl+ and WT dupl- strains on glucose and AMG. (B) Growth curves of WT dupl- cells inoculated as described in the text in 96-well plates containing minimal media with AMG as the sole carbon source (left panel). Subculture of strains to fresh media shows that isolates that began growing between 40 - 60 hours show no lag (right panel). New isolates arise again in some wells. (C) Growth curves of WT dupl+, WT dupl- and six independent mutants of WT dupl- that can grow on AMG as the sole carbon source. In (A) and (C) points and error bars represent the mean and SD of n=3 biological replicates. For each growth curve, the sugar used as the sole carbon source is indicated at the top along with the concentration used.

To better understand mechanisms of regulation of *BT1871-1872* and determine whether duplications or other mutations conferring increased growth on AMG and RFOs could be readily generated de novo, we grew 10 replicates of each of 8 independent colonies of dupl- cells in media with AMG as the sole carbon source. After about 45 hours, we observed growth in 12 wells (Fig. 2B). Upon transfer of the cultures to fresh media containing AMG, dupl- cells from the wells where growth had initially occurred were able to grow with decreased lag times and increased growth rates as compared to the parent dupl- strain, and most isolates grew faster on AMG than dupl+ cells (Fig. 2B, right panel). To confirm that these putative mutants have a stable, heritable phenotype, we saved six independent isolates and tested their growth on glucose, AMG, and melibiose. The isolates grew similar to the parent WT dupl- strain on minimal media with glucose and much better than the parent strain on minimal media with AMG or melibiose (Fig. 2C). Interestingly, the isolates also grew better than WT dupl+ on AMG. Thus, the isolates demonstrate phenotypes that may be caused by a stable genetic change.

To identify the putative mutation(s), we performed long- and short-read whole genome sequencing on three independent isolates and found that each had a different nonsense mutation in *BT1876* (Table S1). *BT1876* encodes the anti-sigma factor paired with the sigma factor *BT1877* in PUL24 (Fig. 3A). To independently test whether inactivation of the BT1876 anti-sigma leads to better growth on RFOs, we made an in-frame deletion in *BT1876* in a WT dupl- background. The *BT1876* mutant strain grew similar to the WT dupl- strain on glucose and substantially better than the WT dupl- strain on AMG and melibiose (Fig. 3B). The *BT1876* strain grew very slightly better than the WT dupl- strain on raffinose and stachyose (Fig. 3B), which both have sucrose moieties that are degraded by other enzymes and do not depend on expression levels of PUL24 genes including *BT1871* (60). Complementation of the *BT1876* mutant with a single copy of *BT1876* in trans under its native promoter restored growth on RFOs to the WT dupl- level (Fig. 3B). Hence, loss of the anti-sigma BT1876 enhances growth of *B. thetaiotaomicron* on RFOs.

**Figure 3.**
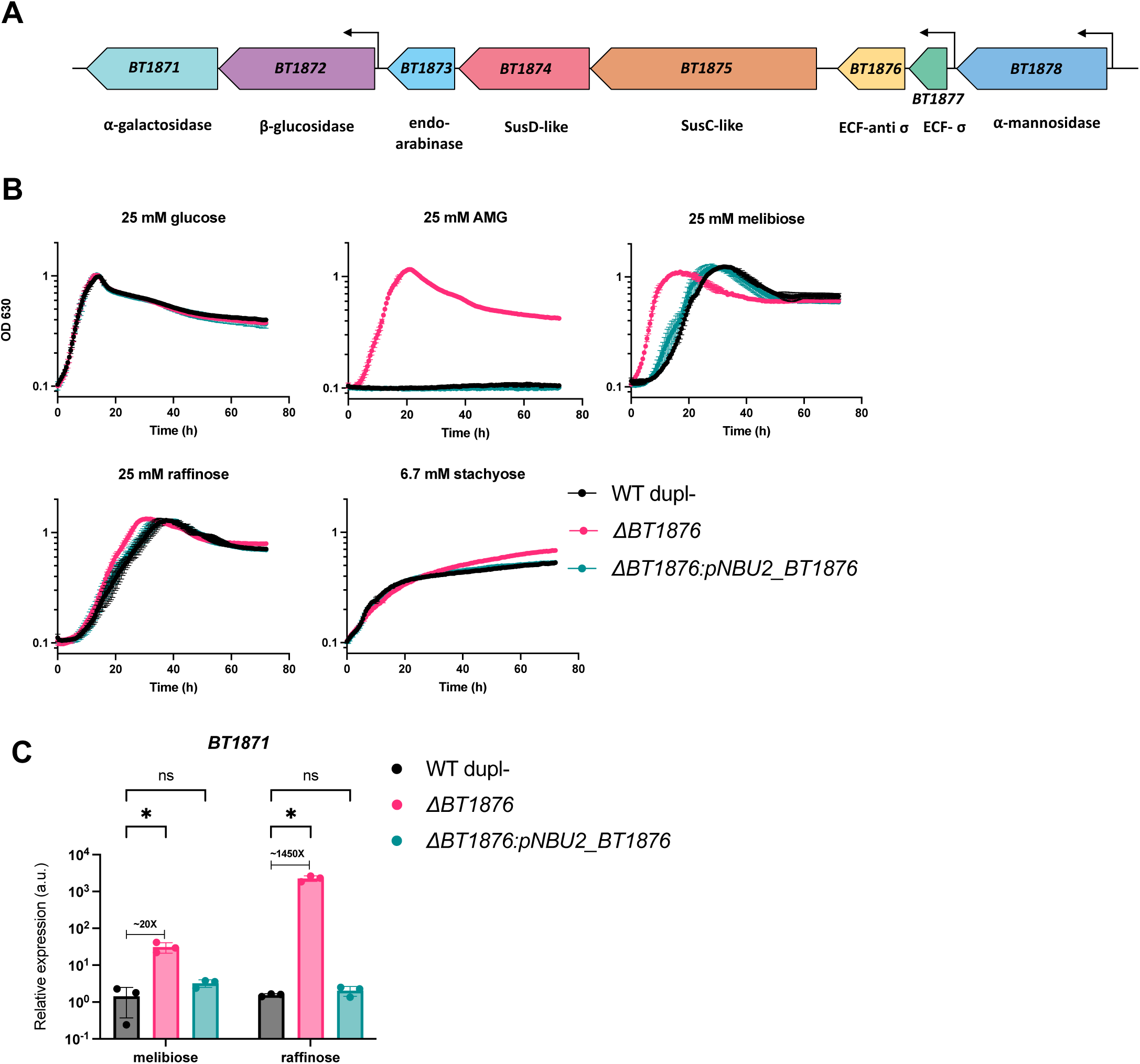
A *BT1876* mutant strain grows better on RFOs than the WT strain due to increased *BT1871* expression. (A) Genomic organization of *B. thetaiotaomicron* PUL24. The location of known transcription start sites ( derived from reference 67 in text) are indicated by bent arrows. (B) Growth curves of WT, *BT1876* anti-sigma mutant and a complemented strain. Points and error bars represent the mean and SD of n=3 biological replicates. For each growth curve, the sugar used as the sole carbon source is indicated at the top along with the concentration used. (C) qRT-PCR showing relative *BT1871* mRNA levels in WT, *BT1876* anti-sigma mutant and the complemented mutant strain. The bars depict mean and SD of n=3 biological replicates. All values are normalized to the WT grown on melibiose. Differences between WT and the *BT1876* mutant are significant based on 2-way ANOVA. *, P < 0.05.

We hypothesized that in the *BT1876* mutant strains, transcription of the α-galactosidase-encoding *BT1871* may be higher than in the WT dupl- strain due to constitutive activity of the sigma factor BT1877. To test this, we used RT-qPCR to measure *BT1871* mRNA levels in WT dupl-, the *BT1876* mutant and complemented mutant strains grown on RFOs as the sole carbon source. We found that *BT1871* expression increased by ∼20-fold on melibiose and ∼1450-fold on raffinose compared to the WT dupl- strain (Fig. 3C). The *BT1871* transcript levels in the complemented strain were similar to the levels in WT dupl- (Fig. 3C).

Two additional lines of evidence confirmed that increased expression of *BT1871* (α-gal) is responsible for better growth of the *BT1876* (anti-sigma) mutant strain on RFOs. First, a *BT1876 BT1871* double mutant strain grew similarly to a *BT1871* single mutant strain on RFOs (Fig. S2A). This suggests that activation of other PUL24 genes in the *BT1876* mutant strain does not confer better growth on RFOs if *BT1871* is absent. Second, a *BT1871* mutant strain is unable to grow on AMG as the sole carbon source and does not yield variants that acquire the ability to grow (Fig. S2B).

Collectively, these data suggest that there are at least two different mechanisms that provide *B. thetaiotaomicron* with increased fitness during growth on RFOs. One mechanism involves duplication of *BT1871* (α-gal) and the other is mutation of *BT1876* (anti-sigma). Both mechanisms result in strongly increased transcription of *BT1871*, encoding an α-galactosidase that breaks the α-1,6 bond of RFOs (60–62). For the remainder of this study, we used a wild-type dupl- strain background to assess other determinants of RFO utilization. For simplicity, we will refer to this strain background as “WT.”

### Other determinants of RFO utilization in *B. thetaiotaomicron*

To determine whether genes outside of PUL24 contribute to RFO utilization, we first tested whether any ⍺-galactosidases other than BT1871 can cleave the ⍺-1,6 bond in RFOs. A recent study showed that when gnotobiotic mice are colonized with a *B. thetaiotaomicron* transposon mutant library, many of the strains that take over the population have transposon insertions upstream of genes encoding ⍺-galactosidases, including *BT1871* (63). The authors hypothesize that increased transcription of the downstream ⍺-galactosidase genes due to readthrough from the transposon gave these mutants an advantage in gut of the mice fed a diet rich in raffinose and melibiose. *BT2851* and *BT3131* were two other genes encoding ⍺-galactosidases for which upstream transposon insertions led to an *in vivo* fitness advantage. To test whether either of these contribute to *B. thetaiotaomicron* RFO use, we constructed mutations in these genes in WT and *BT1871* mutant strain backgrounds. In all cases, strains lacking either *BT2851* or *BT3131* grew similar to their respective parents on RFOs suggesting that these two ⍺- galactosidases do not contribute meaningfully to RFO utilization (Fig. S3A, B).

Full degradation of the RFOs raffinose and stachyose requires an α-galactosidase activity (BT1871 in *B. thetaiotaomicron*) to break the α-1,6 glucose-galactose linkage and at least one other enzyme to break the α-1-β-2 glucose-fructose bond of the sucrose moiety (Fig. S1). *B. thetaiotaomicron BT1871* mutant strains fail to grow on melibiose (which only contains the α-1,6 glucose-galactose linkage) (Fig. S2A), but still grow on raffinose and stachyose, presumably by using fructose liberated by a sucrase that can break the α-1-β-2 glucose-fructose bond. A large-scale functional study found transposon insertions in a *B. thetaiotaomicron* gene cluster (*BT2156-BT2160*) conferred fitness defects on raffinose and a few disaccharides (64).

We constructed *BT2157* and *BT2158* single deletion mutants in both WT and *BT1871* mutant backgrounds to probe their growth on RFOs. Strains with mutations in *BT2157* and *BT2158* grew similar to their respective parent strains on RFOs (Fig. S3C,3D). Thus, *B. thetaiotaomicron* does not use the *BT2156-BT2160* gene cluster to catabolize the sucrose liberated from RFOs.

To identify other genes that may be involved in RFO use, we performed RNA-seq on WT *B. thetaiotaomicron* grown on raffinose or glucose as the sole carbon source. We found 1037 genes significantly differentially expressed (absolute fold change > 2 and adjusted p value < 0.05) on raffinose compared to glucose. Of those, 743 (71.6%) genes were upregulated while 294 (28.4%) were downregulated (Fig. S4 and Table S2). We noted that *BT1871* (α-gal) was not upregulated in raffinose compared to glucose-grown cells even though it confers a growth advantage. We found that genes in PUL22 were highly upregulated (average fold change of ∼776) on raffinose (Fig. S4 and Table S2). *B. thetaiotaomicron* deploys PUL22 for utilizing fructans such as levan and fructooligosaccharides (28). This PUL is induced by fructose through a hybrid two-component sensor kinase BT1754 and contains genes encoding an inner membrane fructose transporter (*BT1758*), a fructokinase (*BT1757*) and three GH32 family enzymes, which include some with sucrase activity (*BT1759*, *BT1760*, *BT1765*) (28). We hypothesized that these PUL22 encoded sucrases cleave the glucose-fructose bond in RFOs, thus liberating free fructose and activating PUL22 through *BT1754*.

To test the role of PUL22 genes in RFO use, we deleted *BT1754,* which encodes the HTCS activator of PUL22. Indeed, a *B. thetaiotaomicron BT1754* mutant strain has a strong growth defect on raffinose and stachyose (Fig. 4A). To determine whether growth on RFOs requires the PUL22 transporter, GH32 sucrases, or both, we constructed *BT1758* (transporter), and *BT1759, BT1760* and *BT1765* (putative sucrases) mutant strains. The *BT1758* (transporter) mutant strain had a growth defect on raffinose and stachyose that was less severe than the growth defect of the *BT1754* (HTCS) mutant strain (Fig. 4A), suggesting that fructose transport via BT1758 is important for growth of *B. thetaiotaomicron* on RFOs. However, mutant strains lacking individual PUL22-encoded sucrases grew as well as the WT parent on raffinose (Fig. 4B).

**Figure 4.**
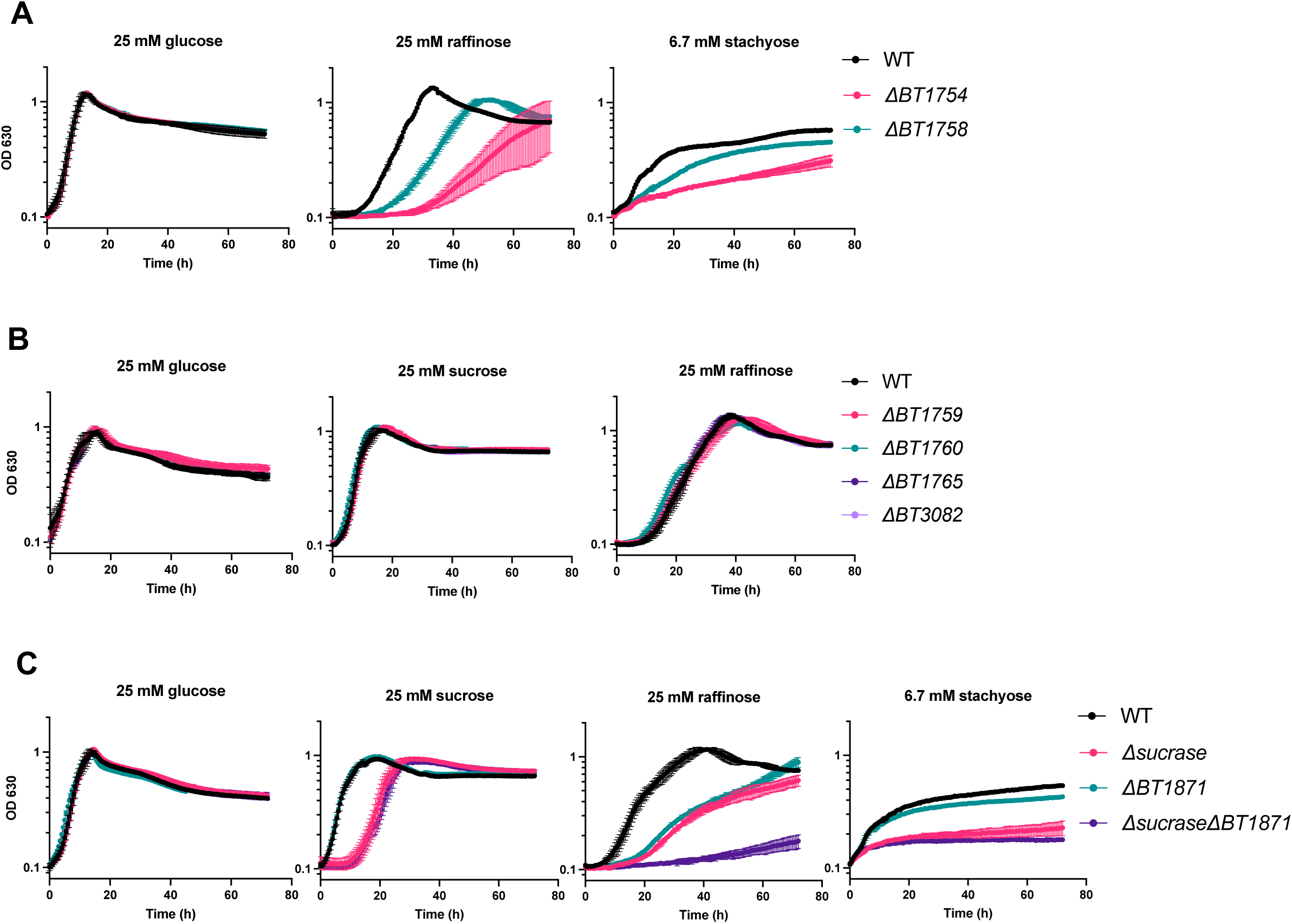
PUL22 is important for RFO utilization in *B. thetaiotaomicron* and its GH32 family sucrases act redundantly to promote RFO utilization. (A) Growth of WT, Δ*BT1754* (sensor kinase) and Δ*BT1758* (putative fructose transporter) on glucose and raffinose. (B) Growth curves of individual GH32 sucrase mutant strains on glucose, sucrose and raffinose. *BT1759, BT1760,* and *BT1765* are part of PUL22. *BT3082* is not part of PUL22 but is coregulated with PUL22 genes in response to fructans. (C) Growth curves of WT, *BT1871* mutant, a quadruple GH32 sucrase mutant (*Δsucrase*) and the *Δsucrase* mutant in a *BT1871* mutant background on RFOs. Points and error bars represent the mean and SD of n=3 biological replicates. For each growth curve, the sugar used as the sole carbon source is indicated at the top along with the concentration.

Another GH32 family enzyme (not encoded in PUL22), BT3082, is produced at high levels in the presence of fructans and has *in vitro* sucrase activity (28). A mutant strain lacking only *BT3082* also had no growth defect on raffinose (Fig. 4B). We then hypothesized that the GH32 family sucrases may act redundantly on RFOs since they all break the glucose-fructose bond. To test this, we constructed a *B. thetaiotaomicron* strain lacking all four GH32 enzymes (Δ*ΒΤ1759* Δ*ΒΤ1760* Δ*BT1765* Δ*BT3082*), which we called the *Δsucrase* mutant strain. This strain grew similar to the WT strain on glucose but had a longer lag phase on sucrose as the sole carbon source (Fig. 4C). Importantly, the *Δsucrase* mutant strain displayed a significant growth defect on raffinose and stachyose compared to the WT strain (Fig. 4C). Combined mutations in *BT1871* (α-gal) and the four GH32 sucrases (*Δsucrase ΔBT1871*) abolished the growth of *B. thetaiotaomicron* on raffinose as the sole carbon source, whereas the *Δsucrase ΔBT1871* and *Δsucrase* strains grew similarly and very poorly on stachyose (Fig. 4C). Taken together, we conclude that *B. thetaiotaomicron* uses enzymes from both PUL22 and PUL24 for utilizing RFOs and the GH32 family sucrases in *B. thetaiotaomicron* have redundant functions with respect to RFO usage.

### RFO dependent activation of PUL24 genes in the *BT1876* anti-sigma mutant requires a global regulator of carbohydrate utilization

We expected that if PUL24 is dedicated to RFO sensing and utilization, the transcription of PUL24 genes would be strongly activated in response to RFOs through inactivation of the anti-sigma BT1876 and release of the sigma factor BT1877. Our RNA-seq analysis suggested only weak activation of PUL24 genes in response to raffinose (Table S2), so we used RT-qPCR to further explore this. We measured levels of two PUL24 mRNAs, *BT1875*, encoding the SusC-like transporter, and *BT1871,* encoding the α-galactosidase important for RFO use. We found that that *BT1875* (transporter) transcripts were upregulated 5.5-fold on melibiose and 4.2-fold on raffinose compared to glucose in the WT strain and *BT1871* (α-gal) was upregulated 3-fold in melibiose and 2-fold in raffinose compared to glucose (Fig. 5A, compare black bars). This level of activation is very small compared to what is seen for activation of other PULs by their cognate substrates. For example, genes in PUL22 are upregulated 50-500-fold in the presence of the cognate substrate levan (28) while genes in PUL7 are upregulated >1000-fold in the presence of the substrate arabinan (34). We reasoned that the fully induced expression levels of PUL24 genes would be observed in the *BT1876* (anti-sigma) mutant strain irrespective of growth substrate if the sigma factor (BT1877) is the primary regulator of PUL24 gene expression. In the *BT1876* (anti-sigma) mutant strain, we observed a 50-fold upregulation of *BT1875* (transporter) compared to the WT strain when grown on glucose (Fig. 5A). Interestingly, levels of *BT1875* (transporter) transcripts in the *BT1876* (anti-sigma) deletion strain were further increased when cells were grown on melibiose (∼700-fold increase compared to WT) and raffinose (∼400,000- fold increase compared to WT). This trend of dramatically increased transcript levels in response to RFOs in the *BT1876* (anti-sigma) mutant background was also observed for *BT1871* (Fig. 5A). These data suggest that there are multiple mechanisms of regulation of PUL24 genes relevant to RFO utilization.

**Figure 5:**
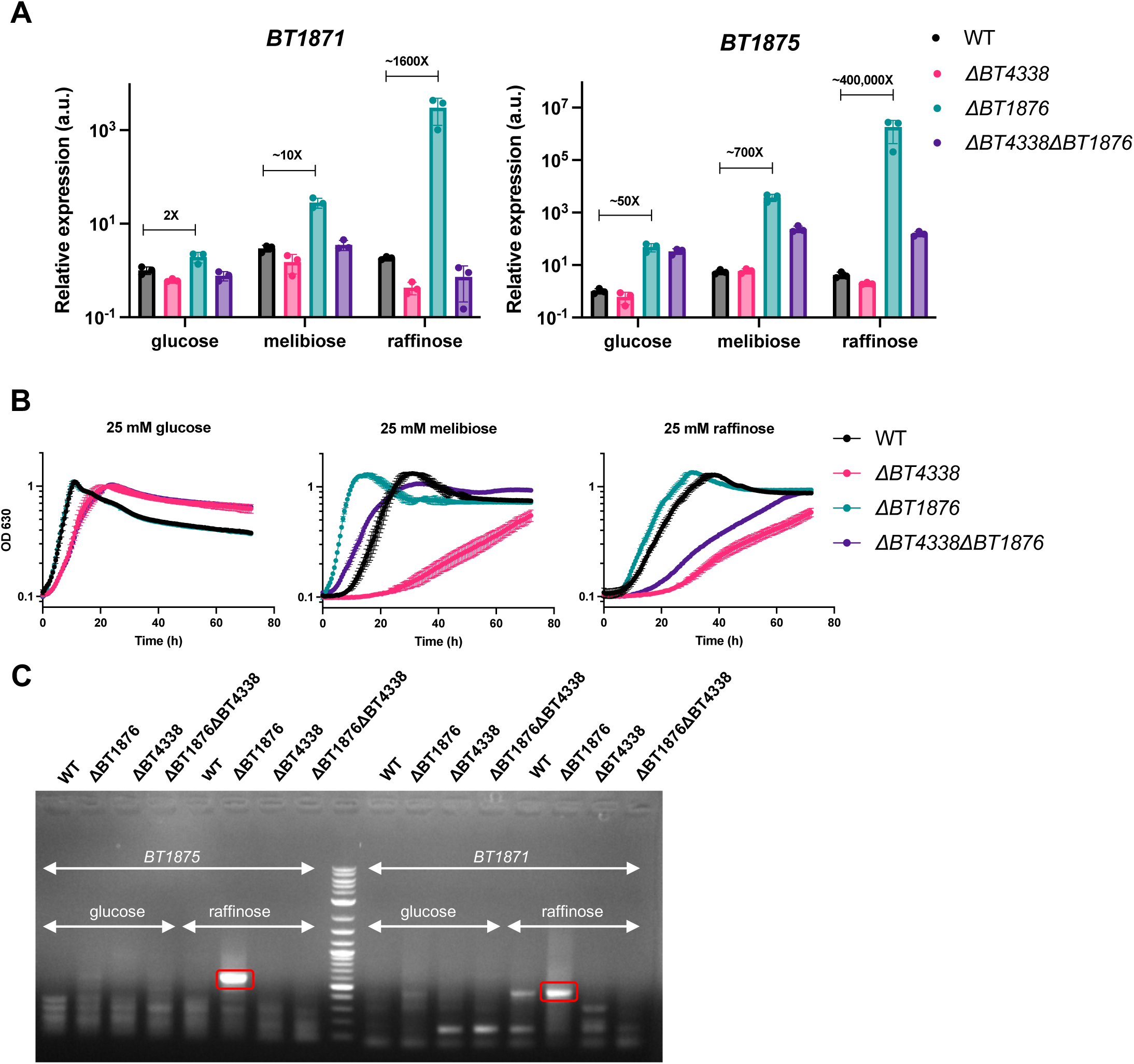
The global regulator *BT4338* is important for RFO utilization in *B. thetaiotaomicron*. (A) qRT-PCR to measure relative levels of *BT1875* and *BT1871* mRNAs. The media and strains used are indicated. The bars depict mean and SD of n=3 biological replicates. All values are normalized to the level in the WT strain grown on glucose. (B) Growth curves of WT, *BT4338* mutant, *BT1876* mutant and their double mutant strains on RFOs. Points and error bars represent the mean and SD of n=3 biological replicates. For each growth curve, the sugar used as the sole carbon source is indicated at the top along with the concentration used. (C) 5’ RACE to identify 5’ ends for *BT1875* and *BT1871* mRNAs. The bands that were sequenced for identifying novel 5’ ends are highlighted in red boxes and the middle lane corresponds to a DNA ladder. Fig. S5 depicts the location of sequenced 5’ ends with respect to the flanking genes.

*B. thetaiotaomicron* possesses a conserved protein BT4338, which has some similarity to the CRP transcription factor that controls the catabolite repression response in enteric bacteria (13, 22, 65). BT4338 is required for efficient use of a variety of monosaccharides and polysaccharides in vitro (34) and many *B. thetaiotaomicron* genes were shown to be differentially regulated in either rich media or under carbon limiting conditions in a *BT4338* mutant strain compared to the wild-type strain (66). We wondered whether BT4338 plays a role in activation of PUL24 genes in response to RFOs. To test this, we measured levels of *BT1875* (transporter) and *BT1871* (α-gal) mRNAs in *BT4338* and *BT4338 BT1876* mutant strains (Fig. 5B). There were small decreases in levels of *BT1875* (transporter) and *BT1871* (α-gal) mRNAs in the *ΔBT4338* mutant strain compared to wild-type in all the conditions tested (Fig. 5A). In the glucose growth condition, Δ*BT1876* (anti-sigma) and Δ*ΒΤ1876* Δ*ΒΤ4338* strains had very similar levels of both the transporter and α-gal transcripts. In sharp contrast, the strong RFO- dependent activation observed for both transcripts in the Δ*BT1876* (anti-sigma) mutant was strongly attenuated in the Δ*ΒΤ1876* Δ*ΒΤ4338* mutant (Fig. 5A). These data suggest that RFO- dependent activation of PUL24 genes in the anti-sigma mutant strain either directly or indirectly depends on BT4338.

We next tested whether growth of these same strains was impacted by the presence or absence of BT4338. There were no major differences in growth when glucose was the sole carbon source between WT, Δ*ΒΤ1876*, Δ*ΒΤ4338,* and Δ*ΒΤ1876* Δ*ΒΤ4338* mutant strains (Fig. 5B). In melibiose, where we know that *BT1871* expression levels are strongly correlated with growth ((60) and Fig. 2B, C), the Δ*ΒΤ4338* strain was strongly growth inhibited compared to the WT parent. This may be explained by the reduced abundance of *BT1871* mRNA observed under this growth condition (Fig. 5A). The growth advantage displayed by the Δ*ΒΤ1876* strain compared to the WT parent was eliminated in the Δ*ΒΤ1876* Δ*ΒΤ4338* strain (Fig. 5B), consistent with reduced *BT1871* mRNA levels observed in the double mutant compared to the Δ*ΒΤ1876* parent (Fig. 5A). In raffinose, where we know that genes in both PUL22 (sucrases) and PUL24 (α-gal) are important for growth, we saw the same overall patterns as for melibiose.

The Δ*ΒΤ4338* strain was strongly growth inhibited compared to the WT parent and the growth advantage displayed by the Δ*ΒΤ1876* strain was eliminated in the Δ*ΒΤ1876* Δ*ΒΤ4338* strain (Fig. 5B). These patterns of growth are consistent with the expression levels of *BT1871* mRNA (Fig. 5A). Taken together, we conclude that *BT4338* is important for RFO utilization in *B. thetaiotaomicron* by modulating (directly or indirectly) transcription of *BT1871* in response to RFOs.

### PUL24 genes are expressed from transcription start sites dependent on BT4338 and BT1877 when grown on RFOs

A recent study annotated transcription start sites (TSSs) in *B. thetaiotaomicron* grown in rich media and identified transcription start sites upstream of *BT1877* and *BT1872* within PUL24 ((67), Fig. 3A). To find putative TSSs in PUL24 that are active during growth on glucose and RFOs, we performed 5’ RACE. We harvested RNA from WT, Δ*BT1876*, Δ*BT4338* and Δ*BT4338* Δ*BT1876* strains grown on glucose or raffinose and used primers to define 5’ ends near *BT1875* and *BT1871.* In the *BT1876* (anti-sigma) mutant grown on raffinose, we observed a strong PCR band representing a 5’ end upstream of *BT1875* (Fig. 5C). A fainter band of the same size was also observed in the *BT1876* mutant grown on glucose. Importantly, this band was absent in the WT, *BT4338* and the *BT4338 BT1876* double mutant on both glucose and raffinose. After sequencing, we mapped this 5’ end to a position 291-nt upstream of the *BT1875* coding sequence (Fig. S5A). In fact, this position is inside the 3’ end of the *BT1876* coding region. We found no consensus promoter sequence for the housekeeping sigma factor upstream of this 5’ end, which suggests that this may represent a site for transcription initiation by the PUL24 sigma factor *BT1877*.

Using primers to detect 5’ ends upstream of *BT1871,* we observed a strong signal in the *BT1876* mutant strain grown on raffinose (Fig. 5C). A fainter band of the same size was also observed for the *BT1876* mutant grown on glucose while there was no band for the *BT4338* mutant or *BT4338 BT1876* double mutant on glucose or raffinose. This signal represented a site 17-nt upstream of the *BT1871* coding sequence (Fig. S5B). These results are consistent with RFO-dependent activation of transcription at a promoter upstream of *BT1871* that is responsive to BT4338.

We hypothesized that *BT1877* (sigma factor) is responsible for both basal (in the WT strain) and activated (in the Δ*BT1876* strain) transcription of PUL24 genes. However, a *BT1877* mutant strain grew similar to the WT strain on RFOs (Fig. S6A), suggesting that *BT1877* is not required for basal transcription of *BT1871*. A Δ*BT1876* Δ*BT1877* double mutant lost the growth advantage of the *BT1876* single mutant when grown on RFOs and completely failed to grow on AMG (Fig. S6B). These observations suggest that the PUL24 sigma factor *BT1877* is not required for basal transcription of *BT1871* but is responsible for its higher transcript levels in the anti-sigma mutant.

Taken together, our results are consistent with a model where activated transcription of PUL24 genes including *BT1871* in response to RFOs requires the PUL24 sigma factor *BT1877* and the global regulator *BT4338*.

### *BT1871* is important for RFO utilization in other *Bacteroides* species

To determine whether mechanisms of RFO utilization are conserved across *Bacteroides* species, we identified homologs of *BT1871* that shared >50% identity across >95% of the protein sequence. To assess the importance these *BT1871* homologs in RFO utilization, we compared the growth of species with *BT1871* homologs namely, *B. thetaiotaomicron*, *Bacteroides ovatus, Bacteroides caccae, Bacteroides faecis, Bacteroides uniformis* and *Bacteroides intestinalis* (Table S3) with growth of species lacking a *BT1871* homolog*, Bacteroides eggerthii*, Bacteroides *fragilis* and *Bacteroides salyersiae*. All these organisms grew similarly on rich (TYG) medium, except *B. eggerthii* which had a long lag phase, while growth in minimal medium with glucose was more variable (Fig. 6A, B). Notably, the group of organisms with a *BT1871* homolog grew on melibiose, albeit with different lags and growth rates (Fig. 6A) while all the species lacking a *BT1871* homolog failed to grow on melibiose as the sole carbon source (Fig. 6B). The two species with the most distant homologs namely, *B. uniformis* and *B. intestinalis* (58% and 56% identity to *BT1871* respectively) had the slowest growth rates and lowest maximum optical densities (OD) when grown on melibiose. This result suggests that having a *BT1871* homolog is important for the growth of *Bacteroides* on melibiose. All the tested strains except *B. eggerthii* DSM 20697 showed some degree of growth on raffinose as the sole carbon source (Fig. 6A, B). Species with *BT1871* homologs tended to have a higher maximal OD on raffinose compared to species without a homolog, although there were no significant differences in raffinose growth rates associated with presence or absence of *BT1871* homolog (Fig. S7A, B). The increased maximal OD may be because the species with *BT1871* homologs can break down the galactose-glucose bond in raffinose as well the glucose-fructose bond thereby using all three monosaccharides whereas species without the *BT1871* homolog are unable to break the galactose-glucose bond and can only use the liberated fructose.

**Figure 6.**
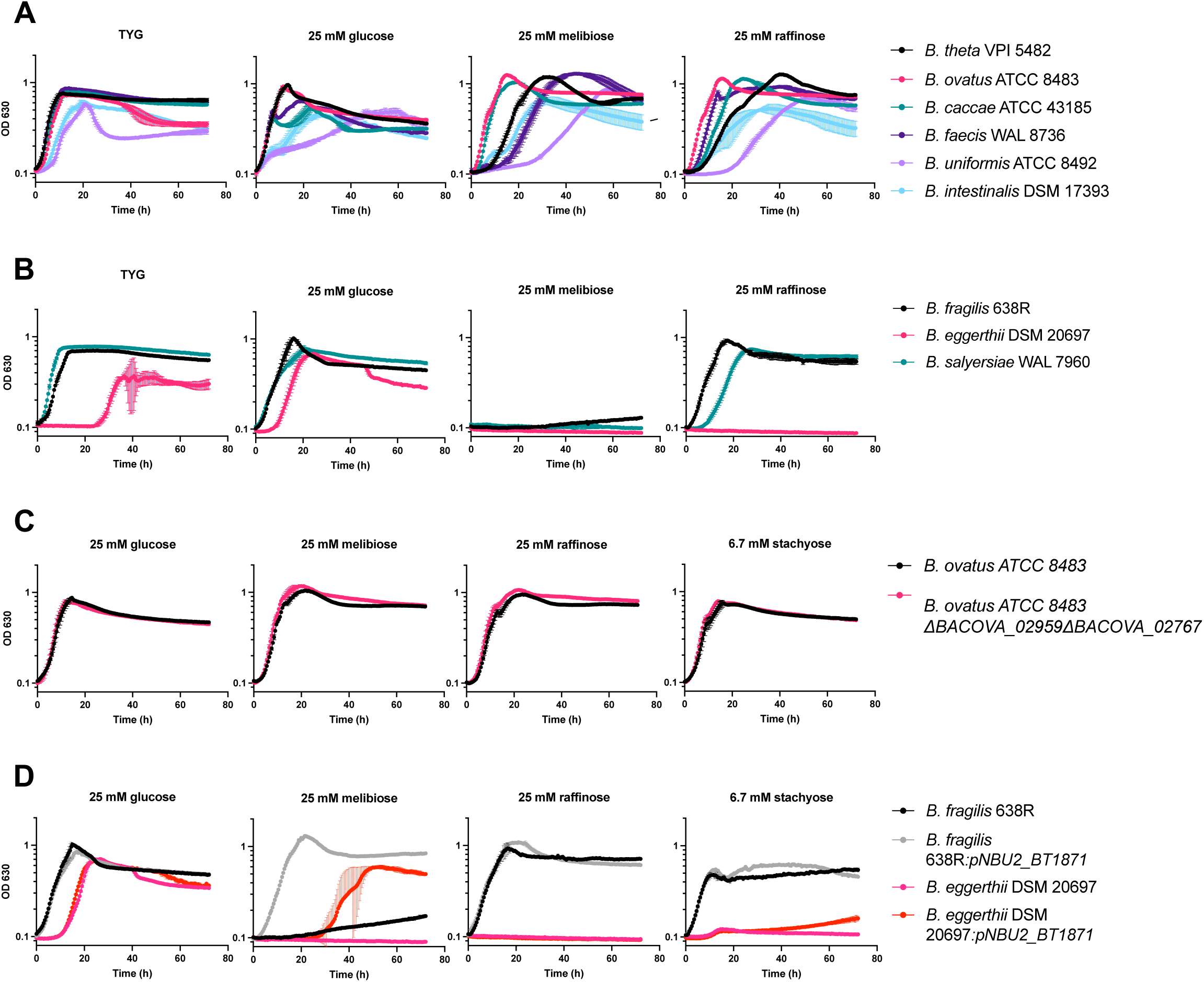
The alpha-galactosidase BT1871 and its homologs are important for melibiose utilization in *Bacteroides* species. (A) Growth curves of *Bacteroides* species containing a *BT1871* homolog on RFOs. (B) Growth curves of *Bacteroides* species lacking a *BT1871* homolog on RFOs. (C) Growth curves of wild-type *B. ovatus* ATCC 8483 (BO) and a mutant strain lacking two *BT1871* homologs on RFOs. (D) Growth curves of *B. fragilis* 638R and *B. eggerthii* DSM 20697 alongside their derived strains expressing *BT1871* from a constitutive promoter. In all panels, points and error bars represent the mean and SD of n=3 biological replicates. For each growth curve, the sugar used as the sole carbon source is indicated at the top along with the concentration used.

To further test the importance of *BT1871* in RFO utilization in other *Bacteroides* species, we deleted the homologs of *BT1871*, namely *BACOVA_02959* and *BACOVA_02767* in *B. ovatus* ATCC 8483. These GH97 family α-galactosidases have 87% and 56% amino acid identity to *BT1871,* respectively (Table S3). Further, *BACOVA_02959* is part of *B. ovatus* ATCC 8483 PUL60 which has synteny to PUL24 in *B. thetaiotaomicron* (32). If these α-galactosidases are necessary and sufficient for cleaving the α-1,6 bond in melibiose, we would expect the double mutant would fail to grow on melibiose as a sole carbon source. However, the double mutant strain did not show a growth defect on RFOs and grew similar to its parent WT strain (Fig. 6C). It is worth noting that *B. ovatus ATCC 8483* has 10 other GH97 family enzymes (with <50% identity to *BT1871*) that might work redundantly to catabolize RFOs (32).

Finally, we wanted to test whether heterologous expression of *BT1871* is sufficient to confer better RFO utilization in organisms that lack a homolog. A constitutive *BT1871* expression cassette was constructed and validated by integration into the genome of a *B. thetaiotaomicron BT1871* mutant strain as a single copy insert. The complemented strain grew better than the *BT1871* mutant and similar to the WT strain on RFOs as the sole carbon source (Fig. S8). This expression cassette was then integrated into the genomes of *B. fragilis 638R* and *B. eggerthii DSM 20697* which lack a *BT1871* homolog. Constitutive *BT1871* expression conferred the ability to grow on melibiose as the sole carbon source on both species (Fig. 6D). However, the *BT1871-*expressing strains grew similar to their WT counterpart when grown on raffinose or stachyose as the sole carbon source (Fig. 6D). Taken together, we conclude that *BT1871* homologs are important for catabolizing melibiose in different *Bacteroides* species.

## Discussion

In this work, we have shown how the model gut commensal *B. thetaiotaomicron* utilizes raffinose family oligosaccharides (Fig. 7). When *B. thetaiotaomicron* encounters RFOs in its environment, they are transported across the outer membrane by an unknown transporter. In the periplasm, the alpha-galactosidase BT1871 cleaves the α-1,6 galactoside bonds, liberating galactose while the remaining sucrose moiety is cleaved by PUL22-encoded GH32 family sucrases. Monomeric fructose activates genes in PUL22 via the HTCS BT1754, resulting in increased expression of these genes. Specific inner membrane proteins then transport the monomers glucose, galactose, and fructose into the cytoplasm where they enter central metabolism. *BT1871* is transcribed at a basal level by an unknown sigma factor and this expression is dependent on the Crp-like protein BT4338. Upon deletion of the anti-sigma *BT1876* or in the presence of the unknown substrate of PUL24, expression of PUL24 is upregulated. The presence of RFOs and BT4338 promote high levels of transcription from novel transcription start sites that are not active under growth in rich media (67). BT4338 may also control expression of PUL22 since CHIP-seq data indicates the presence of several binding sites for BT4338 within PUL22 (66).

**Figure 7.**
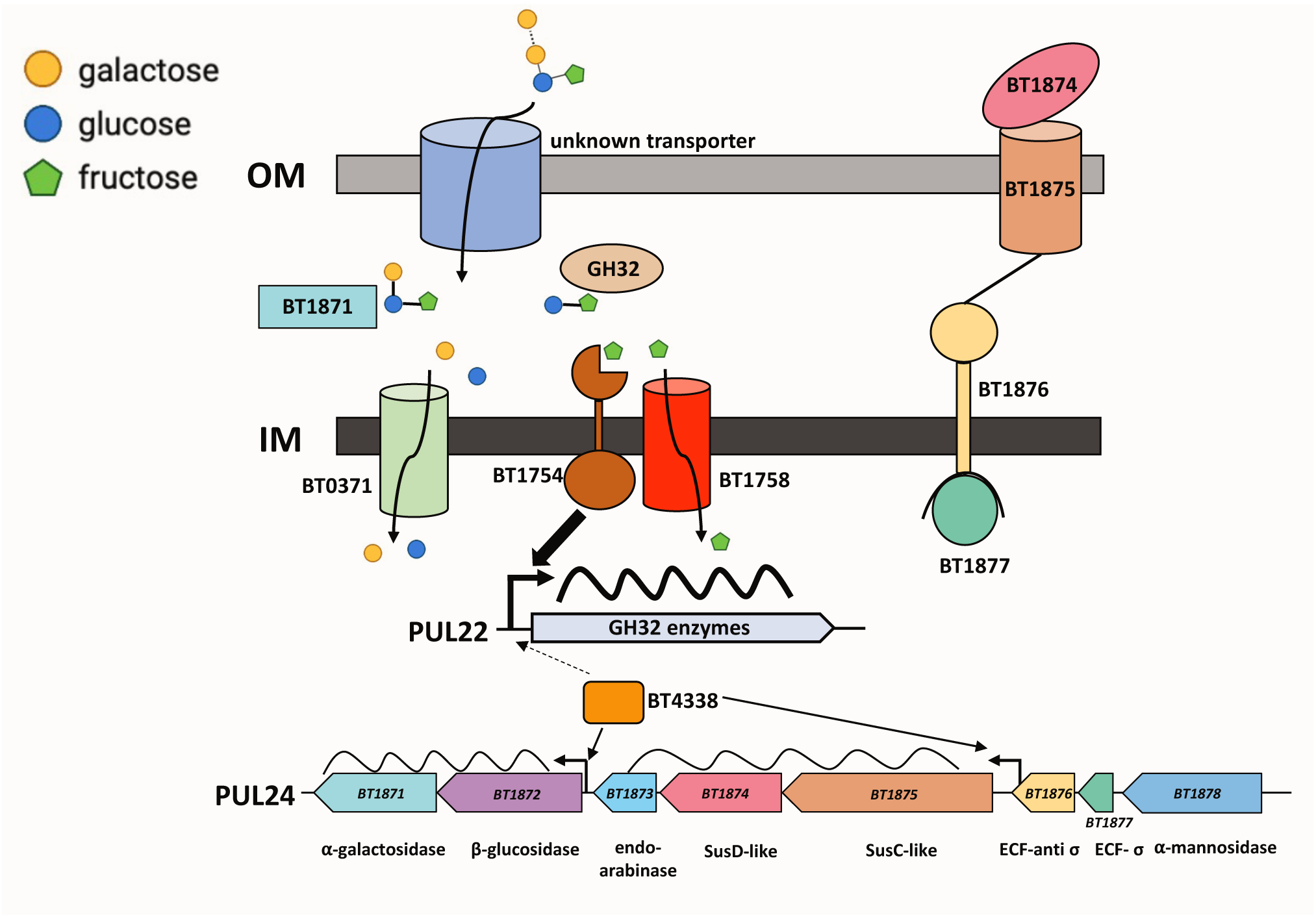
Model for raffinose family oligosaccharide (RFO) utilization by *B. thetaiotaomicron*. After crossing the outer membrane (OM), RFOs are acted upon by BT1871 which cleaves the α-1,6 galactoside bond between glucose and galactose. Next, GH32 family enzymes act redundantly on the liberated sucrose moiety in the periplasm. The monosaccharide subunits are transported across the inner membrane (IM) by dedicated transporters. Expression of PUL22 genes depend on the HTCS BT1754 which binds fructose and activates PUL22 genes, including the sucrases that cleave the bond between glucose and fructose. The natural signal that induces PUL24 genes, including *BT1871*, is unknown, but basal levels of *BT1871* transcription depend on BT4338. When the PUL24 anti-sigma factor is deleted, there is an RFO and BT4338 dependent upregulation from novel transcription start sites in PUL24 whose approximate locations are indicated by bent arrows. A dashed arrow indicates that BT4338 may also control expression of PUL22 genes.

Oligosaccharides are prevalent in a plant rich diet and are gaining popularity as prebiotics (68–70). Here we have shown that hydrolases from two separate PULs, PUL24 and PUL22, work together to degrade an abundant class of plant oligosaccharides, namely RFOs, in *B. thetaiotaomicron*. This dual function of enzymes encoded by PULs capable of degradation of more complex substrates for catabolism of oligosaccharides is also seen for human milk oligosaccharides (HMOs) which are degraded by *B. thetaiotaomicron* using hydrolases from PULs that usually catabolize mucins (71). This is likely due to structural similarity between some components of HMOs and core mucin structures and the presence of similar glycosidic bonds. Similarly, *B. thetaiotaomicron* can utilize galacto-oligosaccharides using enzymes from its pectin-galactan utilization and mucin utilization PULs (72), while fructo-oligosaccharides and arabino-oligosaccharides are degraded using enzymes from the fructan PUL and arabinan PUL, respectively (28, 34). While the natural substrate for PUL24 is still unknown, the presence of an endo-arabinase (*BT1873*) suggests that the substrate could be a plant derived glycan since animal (host) derived glycans do not contain arabinose (73, 74). However, most plant derived glycan degrading PULs contain an HTCS regulator while PUL24 has a sigma, anti-sigma pair controlling it. Further work needs to be done to elucidate the natural substrate of PUL24.

We showed that deleting the master regulator BT4338 affects growth on RFOs and expression of PUL24 genes. However, the mode of action of BT4338 on this PUL is unknown. Does it act as an activator by directly binding to promoters in PUL24 as for *fusA2* (66) or does it indirectly control expression via some other activator as for *roc* (65)? Recent work has shown that BT4338 activated genes are dominantly silenced by the monosaccharides glucose and fructose (65). It is likely that the low level of PUL24 induction in minimal media glucose is because of the same phenomenon and growth on other substrates such as melibiose or raffinose alleviates this silencing. Interestingly, RNA-seq data showed there is transcription from the 3’ region of *BT1876* in nutrient starved *B. thetaiotaomicron* cells while there was no transcription from this region in glucose grown cells (75). The transcription under this condition starts very close to where we mapped the putative 5’ end upstream of *BT1875* using 5’ RACE (Fig. S5A). Importantly, starvation is known to activate *BT4338* and induce widespread gene expression changes in *B. thetaiotaomicron*, possibly by changing levels of some metabolite that modulates BT4338 activity. We failed to find any motif resembling the BT4338 consensus binding sequence ([wwwTATGTTnTAnAACATAwww](http://www.TATGTTnTAnAACATAwww)) (22) upstream of the *BT1875* 5’ end and a CHIP-seq study did not reveal any BT4338 binding sites upstream of *BT1875* (66). This suggests that BT4338 control of *BT1875* expression may be indirect in nature.

Finally, it is important to think about RFO usage by *Bacteroides* in the context of the densely packed and competitive environment of the gut. As mentioned above, *Bacteroides* rely on enzymes encoded by genes in PULs for RFO degradation. However, other gut commensals like *Bifidobacterium* and *Lactobacillus* harbor operons that encode dedicated RFO regulation, transport, and metabolism functions (51, 53, 57). This may reflect that *Bacteroides* are better adapted to preferentially degrade longer polysaccharides whereas other commensals are more efficient degraders of smaller oligosaccharides like RFOs or HMOs. Indeed, *Bifidobacterium animalis* subsp. *lactis Bl-04*, which has a dedicated RFO utilization system (57), can outcompete *B. ovatus* when grown on raffinose as the sole carbon source. Moreover, *in vitro* fermentation of fecal inoculum containing raffinose from healthy human donors led to increased abundance of *Bifidobacteriales* and *Lactobacillales* with a concurrent decrease in *Bacteriodales* (54). These observations suggest that in microbial communities where *Bacteroides* species are competing with organisms with dedicated RFO utilization systems, *Bacteroides* may not be the primary RFO users. However, a recent study showed that acquisition of mutations like the ones we report here may indeed be relevant *in vivo* for *Bacteroides* RFO utilization. The authors of this study found that expression of genes in PUL24 increased when germ-free mice fed an RFO-rich diet were monocolonized with *B. thetaiotaomicron* (63), and they found evidence of spontaneous duplication of the *BT1871* locus in these mice. Strains with *BT1871* duplications in their study varied with respect to the length of the duplicated region and exact junctions, but all showed the same outcome as we observed: increased expression of *BT1871* due to transcription from a ribosomal promoter driving the duplicated copy. These studies were performed in mice fed a diet rich in melibiose and raffinose where increased expression of *BT1871* would be advantageous. Future functional studies using different diets and competition assays with other gut commensals will be needed to fully elucidate the importance of PUL24 and RFO utilization for *Bacteroides in vivo*.

## Materials and Methods

### Media and growth conditions

All strains, plasmids, and oligonucleotides used in this study are listed in Table S4, S5 and S6 respectively. *B. thetaiotaomicron* VPI-5482 and other *Bacteroides* strains were grown in a Coy Laboratory Products vinyl anaerobic chamber with an input gas of 20% CO_2_, 10% H_2_, and 70% N_2_ balance. Routine culturing of *Bacteroides* was done in TYG (tryptone, yeast extract, and glucose) broth (76) and on Difco brain heart infusion (BHI) agar plates supplemented with 10% defibrinated horse blood (HB; Quad Five) or 5 mg/liter hemin and 2.5 μg/liter vitamin K3 (BHIS) at 37°C. *Escherichia coli* strains were grown aerobically at 37°C on BHIS for conjugations and in LB for all other applications. Minimal medium (77) was supplemented with B_12_ (3.75 nM, final; Sigma) and carbohydrates as needed at a final concentration of 25 mM except stachyose which was used at 0.5% w/v. When needed, antibiotics were added at the following final concentrations: 100 μg/ml ampicillin (Sigma), 200 μg/ml gentamicin (Goldbio) and 25 μg/ml erythromycin (VWR).

### Construction of strains and genetic manipulation

In-frame gene deletions and mutations were generated using the counter-selectable allelic exchange vector pLGB13 (78). Flanking regions of 1 kb on either side of the target gene were PCR amplified using Q5 polymerase (NEB) and cloned into pLGB13 using the NEBuilder HiFi cloning kit (NEB). The pLGB13 plasmid was linearized by double digest with BamHI and SalI restriction enzymes. Ligated plasmids were transformed into *E. coli* S-17 Lambda pir cells and positive clones were identified using colony PCR (Gotaq green) and sent for whole plasmid sequencing (Plasmidsaurus). Overnight cultures of the recipient *Bacteroides* as well the donor *E. coli* strains were diluted 1:100 and 1:200 respectively in 5 ml fresh medium and grown for ∼6 h. 1 ml of each culture was centrifuged at 5000 x g for 10 mins and both pellets were combined by resuspending in 100 ul 1X PBS. The suspension was spread out on BHIS plates and incubated aerobically at 37° C overnight to form a lawn. The next day, the lawn was scraped and resuspended in 5 ml 1X PBS. 5 μl of this suspension was spread on BHIS with erythromycin and gentamicin plates and incubated anaerobically. After 2-3 days, 2 colonies were selected and restreaked for isolation and colony PCR was performed to verify cointegrants. For colony PCR, a single colony was lysed for 15 minutes at 95° C in 20 μl of 25 mM NaOH, 0.2 mM disodium EDTA, pH 12. The lysate was neutralized with 40 μl of 40 mM Tris-HCl, pH 5.0 which was diluted 1:2 in ddH_2_O. 2 μl of the diluted lysate was used for subsequent PCR. For counter selection, a single cointegrate colony was grown overnight in TYG, diluted 1:100 in 5 ml fresh TYG and grown for ∼ 6h. 3 μl of the culture was spread on BHIS plates containing 100 ng/ml of anhydrotetracycline (Sigma) for counter selection. After 3-4 days, 8 colonies were inoculated into 200 μl TYG in a 96 well plate and grown overnight. 4 μl of the cultures were lysed as mentioned above and analyzed by PCR to screen for gene deletion.

Complementation of deletion strains and introduction of *BT1871* in other *Bacteroides* species was done using the pNBU2-ErmG vector (79). The gene to be introduced was cloned into the pNBU2 vector, transformed into *E. coli* S-17 cells and positive clones identified. The vector was conjugated into recipient *Bacteroides* strains as described above. After conjugation, cells were spread on BHIS with erythromycin and gentamycin plates. After 2-3 days, two colonies were restreaked on BHIS plus erythromycin plates for isolation. A single integrant was selected, grown overnight in TYG with erythromycin and 4 μl of the culture was lysed as above and subjected to PCR to verify which *att* site the plasmid has integrated into. A further PCR verification was done using primers specific to pNBU2 to confirm integration into the genome.

### Minimal media carbon growth assays

Strains were cultured in triplicate in 5 ml TYG overnight to stationary phase. 1 ml of overnight cultures was spun down at 5,000 × g for 10 min at room temperature. Supernatants were removed, and the pellets were washed twice by resuspending in 1 ml of minimal medium without a carbon source. Next, 2 μl of resuspended cells was inoculated into 198 μl of minimal media containing carbon sources in flat-bottom, 96-well Corning Costar tissue culture-treated plates (catalog #3598). Plates were sealed with a Breathe-Easy gas permeable membrane (Sigma) and statically cultured in the BioTek plate reader at 37° C for as long as needed (usually 72 h) with the optical density recorded every 30 min. Growth rates were measure by first manually inspecting the individual growth curves and removing points that fall in the stationary phase and then fitting the growth curves to a logistic curve.

### Measurement of gene expression by RT-qPCR

Strains were grown in triplicate in 5 ml TYG to stationary phase overnight and culture volumes equivalent to 2 OD_600_ units were washed twice by centrifuging at 5,000 × g for 10 mins and resuspending in 1 ml minimal medium without a carbon source. Washed cells were inoculated 1:100 in 5 ml minimal medium containing a carbon source. All cultures for strains containing pNBU2_ermG vectors contained erythromycin. Cultures were grown to mid-exponential phase at an OD_600_ ∼0.8. Next, 5 ml of cells were pelleted at 4,000 × g for 10 min at 4° C, supernatant decanted, and then RNA was isolated with a Qiagen RNeasy minikit according to manufacturer’s instructions. Residual DNA was degraded using 4-5 μl DNase-I (Thermo) and RNA was cleaned up using phenol-chloroform followed by an overnight ethanol, sodium acetate precipitation and quantified using qubit assay kit (Thermo) or nanodrop. We used a probe-based approach for RT-qPCR to reduce off target amplification and allow quantification of target genes and normalization (using 16S rRNA) in a single reaction. cDNA synthesis and PCR amplification was done with the Luna® Probe One-Step RT-qPCR 4X Mix with UDG according to manufacturer’s instructions. 10 μl reactions were performed using 20-25 ng RNA as input. 400 nM of gene specific primers and 200 nM of gene specific probes were used. The probe for the 16S rRNA gene was labeled with 5’-FAM (IDT) while *BT1871* and *BT1875* specific probes were labeled with 5’-HEX (IDT). Using the ddCT method, raw values were normalized to 16S rRNA values and then minimal medium with melibiose or raffinose values were referenced to the values obtained in minimal medium with glucose to obtain a fold-change.

### 5’ RACE

5’ RACE was done using a Template Switching (TS) Reverse Transcriptase Enzyme mix (NEB) that takes advantage of a template switching reverse transcriptase and a template switching oligonucleotide (TSO). The protocol is identical to the one in (80). Briefly, 250 ng of total RNA (in 4 μl) was mixed with 1 μl random hexamers (50 μM), 1 μl dNTP (10 mM) and heated at 70° C for 5 mins and kept on ice. 2.5 μl of TS buffer (4X), 0.5 μl of a template switching oligo (75 μM) and 1 μl of TS Enzyme mix (10X) were added to the RNA and incubated at 42° C for 90 min and then at 85 ° C for 5 mins. The RT reaction was diluted 2-fold with nuclease free water and 1 μl of the diluted mix was subjected to 5’ RACE PCR using Q5 polymerase (NEB) with a touchdown PCR protocol. A TSO specific primer and a gene specific primer were used for each PCR reaction. The PCR products were separated on a 1% agarose gel and bands were excised using a QIAquick gel extraction kit (Qiagen). The purified PCR bands were cloned using the NEB PCR cloning kit following manufacturer’s instructions and transformed colonies were screened using colony PCR. Positive colonies were grown overnight, plasmids extracted and sent for whole plasmid sequencing.

### Whole genome sequencing sample preparation, processing and analysis

Overnight cultures were grown, pelleted, and total high molecular weight genomic DNA was purified using a DNEasy blood and tissue kit (Qiagen) according to manufactures instructions. A total of 500 μl of overnight culture was used to extract genomic DNA. Purified DNA was submitted for whole genome sequencing at SeqCenter (Pittsburgh, PA) using both Illumina and Oxford Nanopore sequencing. For the WT dupl- mutants capable of growth on AMG as the sole carbon source, sample libraries for Illumina sequencing were prepared using the Illumina DNA Prep kit and IDT 10bp unique dual indices indices, and sequenced on an Illumina NextSeq 2000, producing 2×151bp reads. Demultiplexing, quality control and adapter trimming was performed with bcl-convert (v3.9.3) (81). Nanopore samples were prepared for sequencing using Oxford Nanopore’s “Genomic DNA by Ligation” kit (SQK-LSK109) and protocol. All samples were run on Nanopore R9 flow cells (R9.4.1) on a MinION. Guppy (v5.0.16) (82) was used for high-accuracy basecalling, demultiplexing, and adapter removal. Porechop (v0.2.3_seqan2.1.1, default parameters) (83) was used to trim residual adapter sequence from Nanopore reads that may have been missed during basecalling and demultiplexing. Hybrid assembly with Illumina and Oxford Nanopore reads was performed with Unicycler (v0.4.8, default parameters) (84). Assembly statistics were recorded with QUAST (v5.0.2, default parameters) (85) and annotation was performed with Prokka (v1.14.5, default parameters + ‘-- rfam’) (86)p. For WT dupl+, WT dupl-, Δ*rbpB* dupl+ and Δ*rbpB* dupl- genomes, sample libraries for Illumina sequencing were prepared using the tagmentation-based and PCR-based Illumina DNA Prep kit and custom IDT 10bp unique dual indices with a target insert size of 280 bp.

Illumina sequencing was performed on an Illumina NovaSeq X Plus sequencer producing 2×151bp paired-end reads. Demultiplexing, quality control and adapter trimming was performed with bcl-convert (v4.2.4). For Nanopore sequencing, sample libraries were prepared using the PCR-free Oxford Nanopore Technologies (ONT) Ligation Sequencing Kit (SQK-NBD114.24) with the NEBNext Companion Module (E7180L) to manufacturer’s specifications. Nanopore sequencing was performed on an Oxford Nanopore a MinION Mk1B sequencer or a GridION sequencer using R10.4.1 flow cells in one or more multiplexed shared-flow-cell runs. Run design utilized the 400bps sequencing mode with a minimum read length of 200bp. Guppy (v6.4.6) (82)was used for super-accurate basecalling (SUP), demultiplexing, and adapter removal. Porechop (v0.2.4, default parameters) (83) was used to trim residual adapter sequences from Nanopore reads that may have been missed during basecalling and demultiplexing. *De novo* genome assemblies were generated from the Oxford Nanopore Technologies (ONT) read data with Flye2 (v2.9.2, --asm-coverage 50, --genome-size 6000000) (87) under the nano-hq (ONT high-quality reads) model. Subsequent polishing used the Illumina read data with Pilon (v.1.24, default parameters) (88). To reduce erroneous assembly artifacts caused by low quality nanopore reads, long read contigs with an average short read coverage of 15x or less were removed from the assembly. Assembled contigs were evaluated for circularization via circulator (v1.5.5) (89) using the ONT long reads. Assembly annotation was then performed with Bakta (v1.8.1) (90) using the Bakta v5 database. Finally, assembly statistics were recorded with QUAST (v5.2.0) (85). For the AMG mutants, short-read mapping for mutation detection was completed with breseq (v0.33.2, default parameters) (91). DNA-seq processing statistics are summarized in Table S7.

### RNA sequencing sample preparation, processing, and analysis

Strains were grown in triplicate in 5 ml TYG to stationary phase overnight and cultures equivalent to 2 OD_600_ units were washed twice by centrifuging at 5,000 × g for 10 mins and resuspending in 1 ml minimal medium without a carbon source. Washed cells were inoculated 1:100 in 5 ml minimal medium containing glucose or raffinose to a final concentration of 25 mM. Total RNA was isolated from 5 ml cultures and quantified as described above. 1 μg of total RNA was then submitted to SeqCenter for rRNA depletion, library construction, and sequencing.

Briefly, samples were DNAse treated with Invitrogen DNAse (RNAse free). Library preparation was performed using Illumina’s Stranded Total RNA Prep Ligation with Ribo-Zero Plus kit and 10bp unique dual indices. Sequencing was done on a NovaSeq 6000, producing paired end 151bp reads. Demultiplexing, quality control, and adapter trimming was performed with bcl-convert (v4.1.5). Fastq files were quality checked using FastQC (v0.11.9) (92) and aligned to the genome of *B. thetaiotaomicron* VPI-5482 type strain (ASM1106v1) using Bowtie2 (v2.5.0, default parameters) (93). Aligned reads were quantified using FeatureCounts (v2.0.3, paired end, -s parameter set to 2) (94). Differential expression analysis was done in Rstudio (v2023.12.0+369) using DESeq2 (v1.44.0) (95) with α (False Discovery Rate) set to 0.05 and cutoffs for differentially expressed genes set to absolute value of log 2 fold change >1 and adjusted p value < 0.05.

RNA-seq processing statistics are summarized in Table S7.

### Figures and statistical analysis

All growth curves were plotted, and statistical analyses done using Prism (version 10.0.3). PowerPoint and Biorender were used to make figures and models.

## Data availability

DNA and RNA sequencing datasets from this manuscript are publicly available from NCBI under BioProject accession number PRJNA1107238.

## Acknowledgements

We thank the members of AB’s thesis committee, James Imlay, Bill Metcalf, and Shannon Sirk, for their advice throughout the course of this project. We also thank the former and current Vanderpool and Slauch laboratory members and members of the Microbiome Metabolic Engineering theme for thought-provoking discussions. This work was funded by National Institutes of Health grant R01 R35 GM139557 to CKV. This research was also supported by the GEMS Biology Integration Institute, funded by the National Science Foundation DBI Biology Integration Institutes Program, Award #2022049 (CKV).

## Supplementary Figures

**Figure S1.**
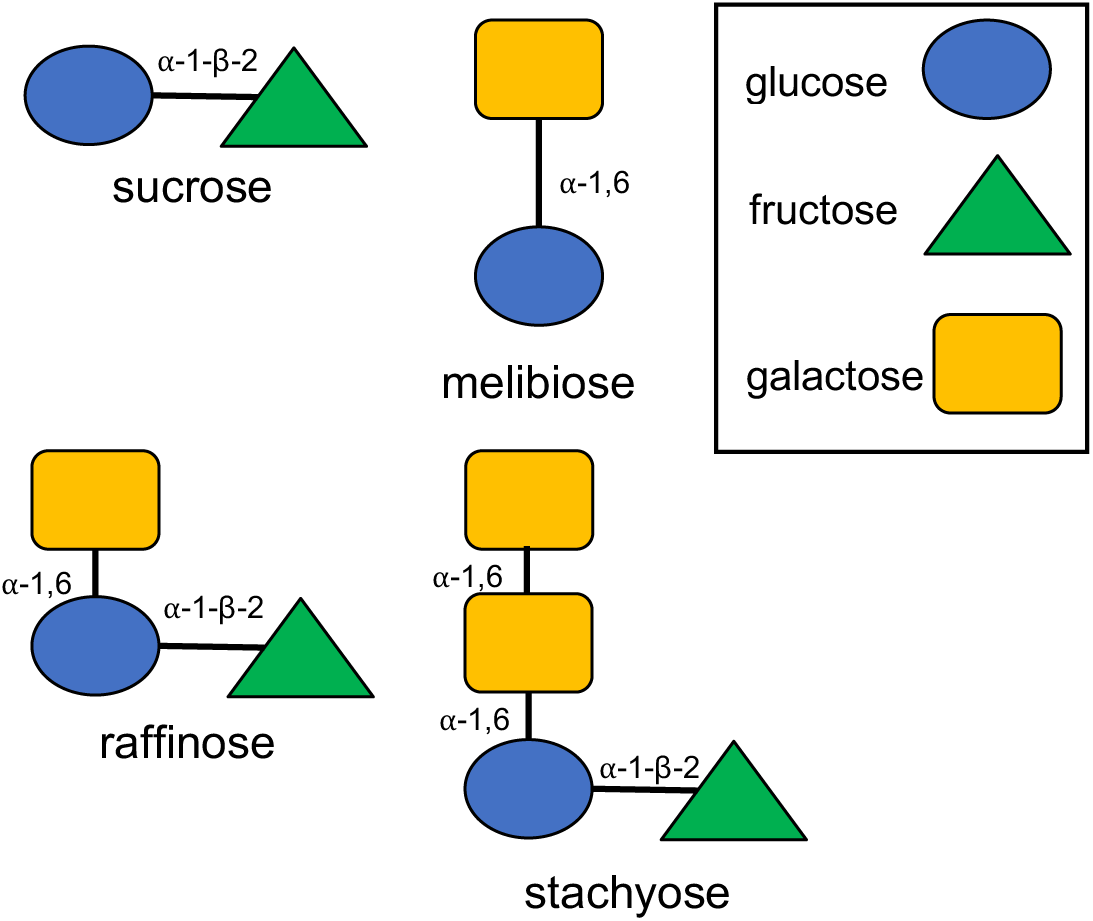
General structure of Raffinose Family Oligosaccharides (RFOs). RFOs are represented with subunits and their linkages indicated next to them. Sucrose and melibiose are disaccharide subunits of RFOs. The monosaccharide subunits which make up RFOs are indicated in the box.

**Figure S2.**
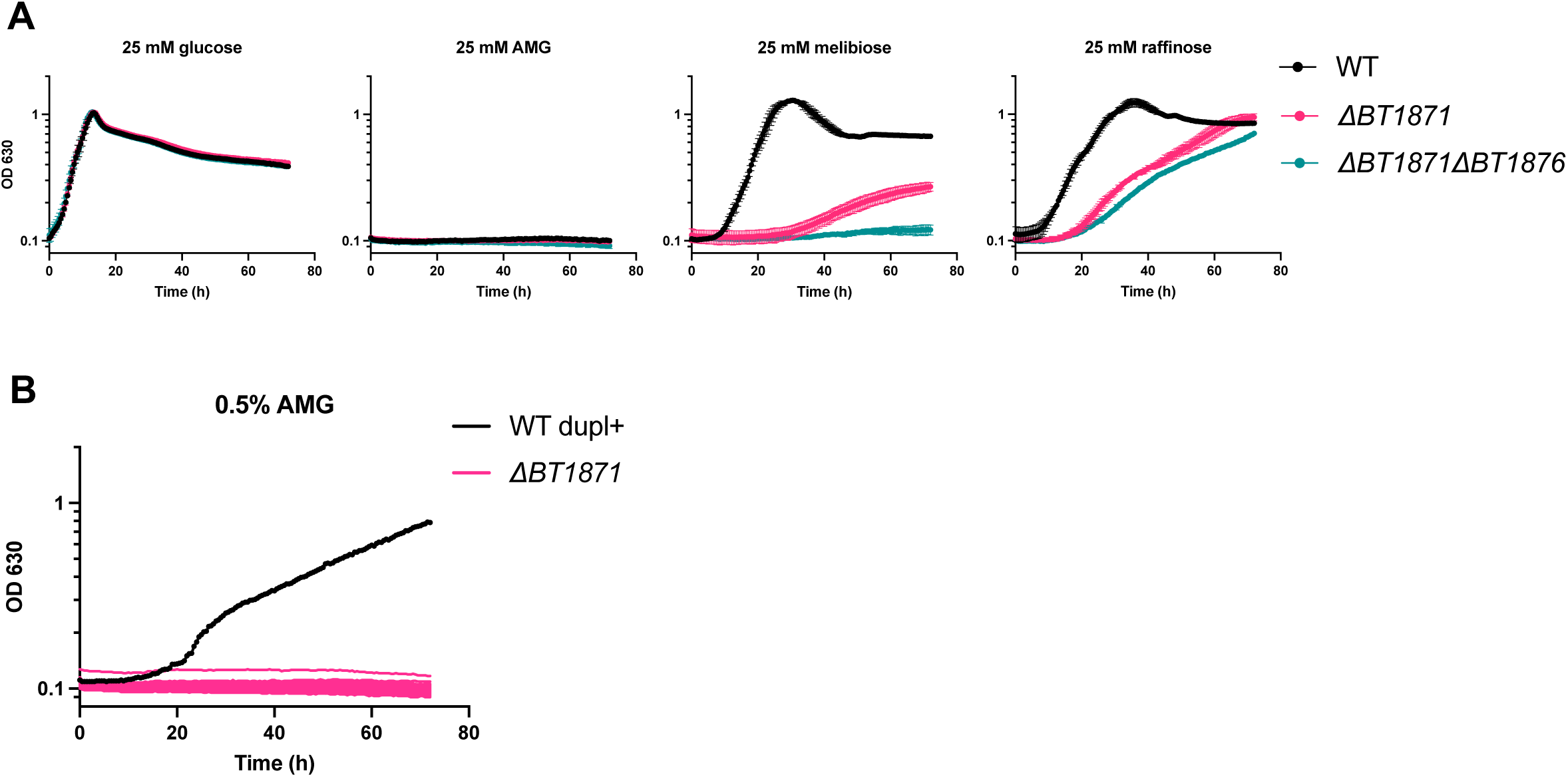
*BT1871* is the only gene in PUL24 important for RFO utilization in *B. thetaiotaomicron*. (A) Growth curves of WT dupl-, *BT1871* mutant and *BT1871 BT1876* double mutant strains on RFOs. Points and error bars represent the mean and SD of n=3 biological replicates. (B) One representative growth curve from an assay to find mutants capable of growth on AMG as in Fig. 2C. Growth curves from individual wells are depicted. For each growth curve, the sugar used as the sole carbon source is indicated at the top along with the concentration used.

**Figure S3.**
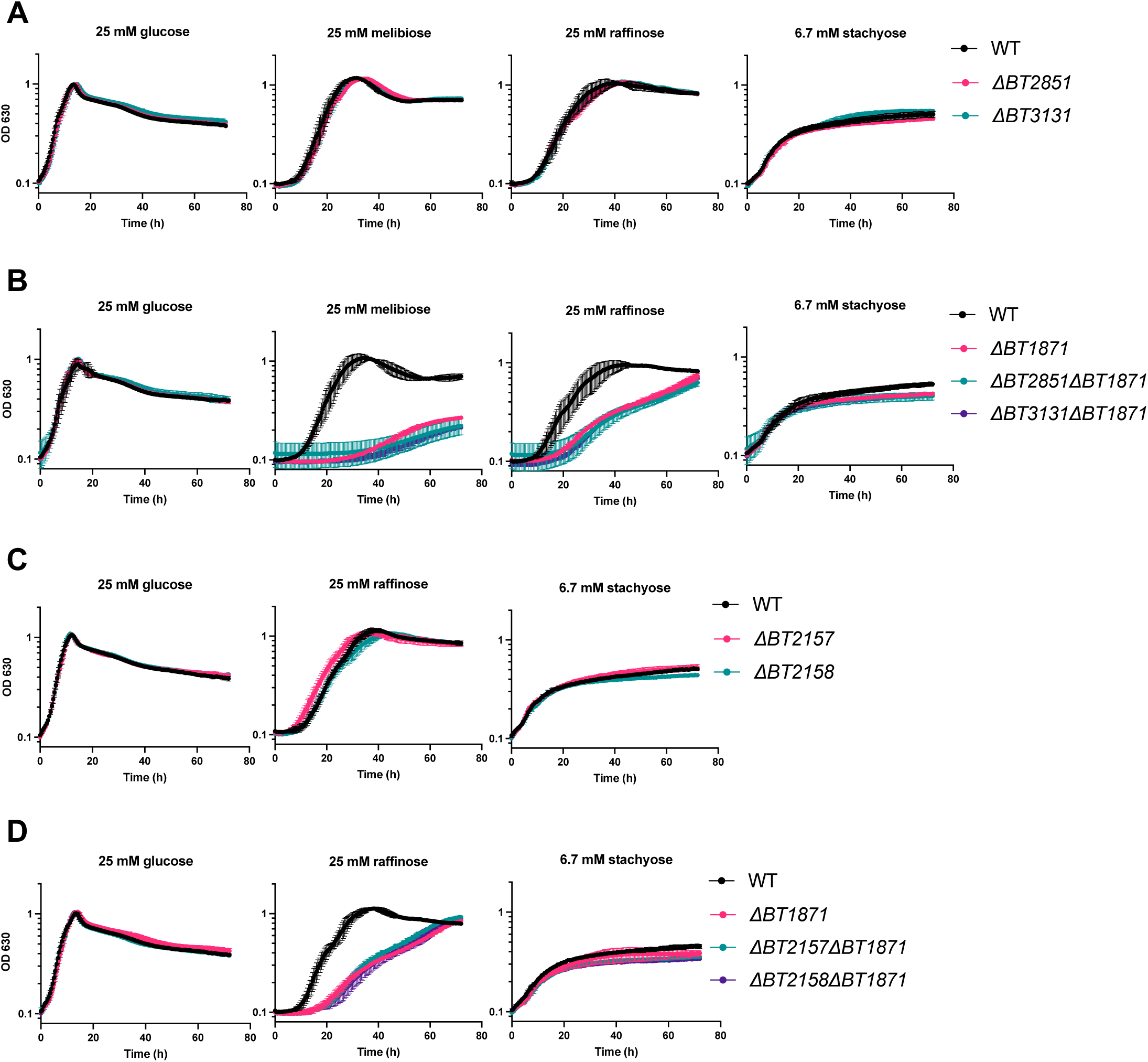
The ⍺-galactosidases *BT2851* and *BT3131,* the dehydrogenase *BT2158,* and the glycosidase *BT2157* are not involved in RFO utilization in *B. thetaiotaomicron*. (A) Growth curves of WT, *BT3131,* and *BT2851* mutant strains on RFOs. (B) Growth curves of WT, *BT1871*, *BT2851 BT1871,* and *BT3131 BT1871* mutant strains on RFOs. (C) Growth curves of WT, *BT2157,* and *BT2158* mutant strains on RFOs. (D) Growth curves of WT, *BT1871*, *BT1871 BT2157*, *BT1871 BT2158* mutants on RFOs. In all panels, points and error bars represent the mean and SD of n=3 biological replicates. For each growth curve, the sugar used as the sole carbon source is indicated at the top along with the concentration used.

**Figure S4.**
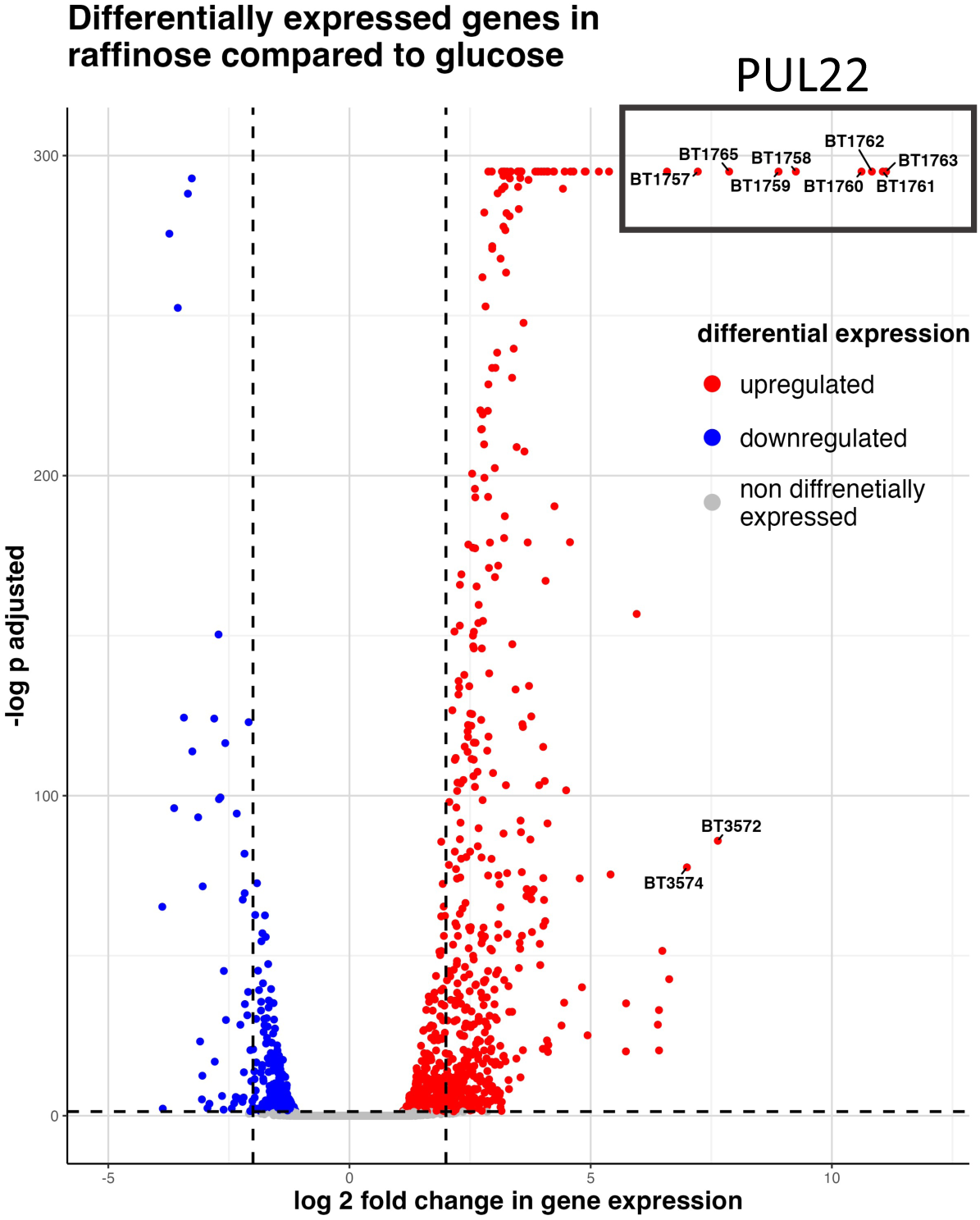
Growth of *B. thetaiotaomicron* on raffinose as the sole carbon source leads to upregulation of PUL22 genes compared to growth on glucose. Volcano plot depicting differentially expressed genes on raffinose compared to glucose. The dashed black lines indicate the threshold used to determine significance (absolute log 2 fold change > 1 and adjusted p value < 0.05). Top 10 highest differentially expressed genes are labeled. The black box highlights PUL22 genes which are highly upregulated on raffinose.

**Figure S5.**
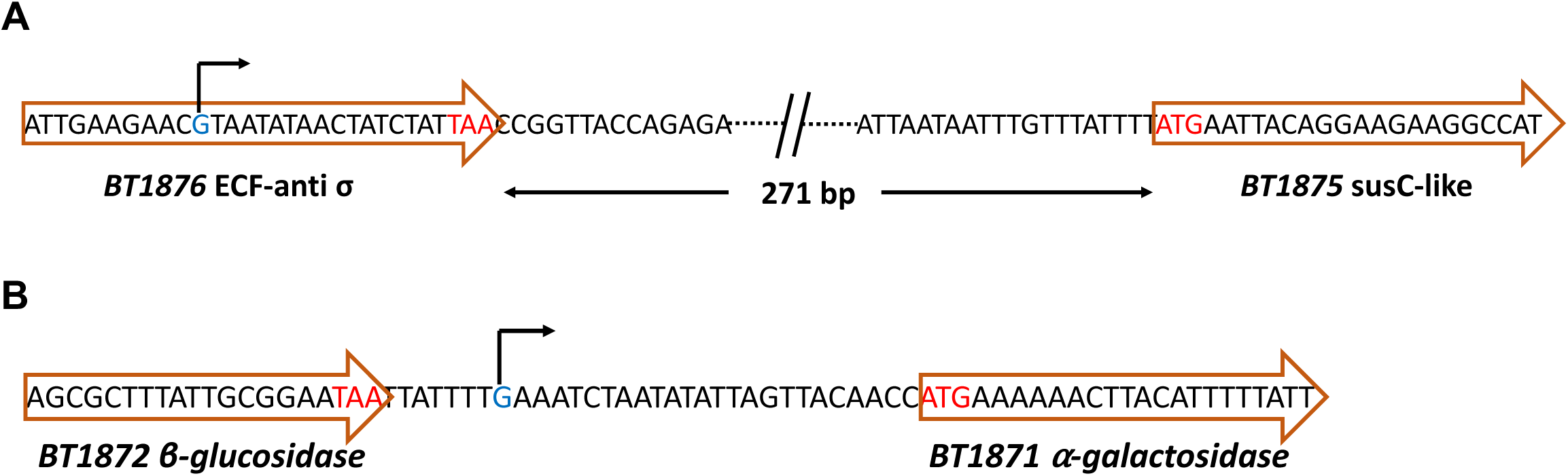
Locations of novel 5’ ends found in PUL24 using 5’ RACE. (A) and (B) represent the location of novel 5’ ends for *BT1875* and *BT1871* respectively, annotated using 5’ RACE as depicted in Fig. 5C. The identified ends are shaded blue with a bent arrow on top. The start and stop codons of genes are shaded red.

**Figure S6.**
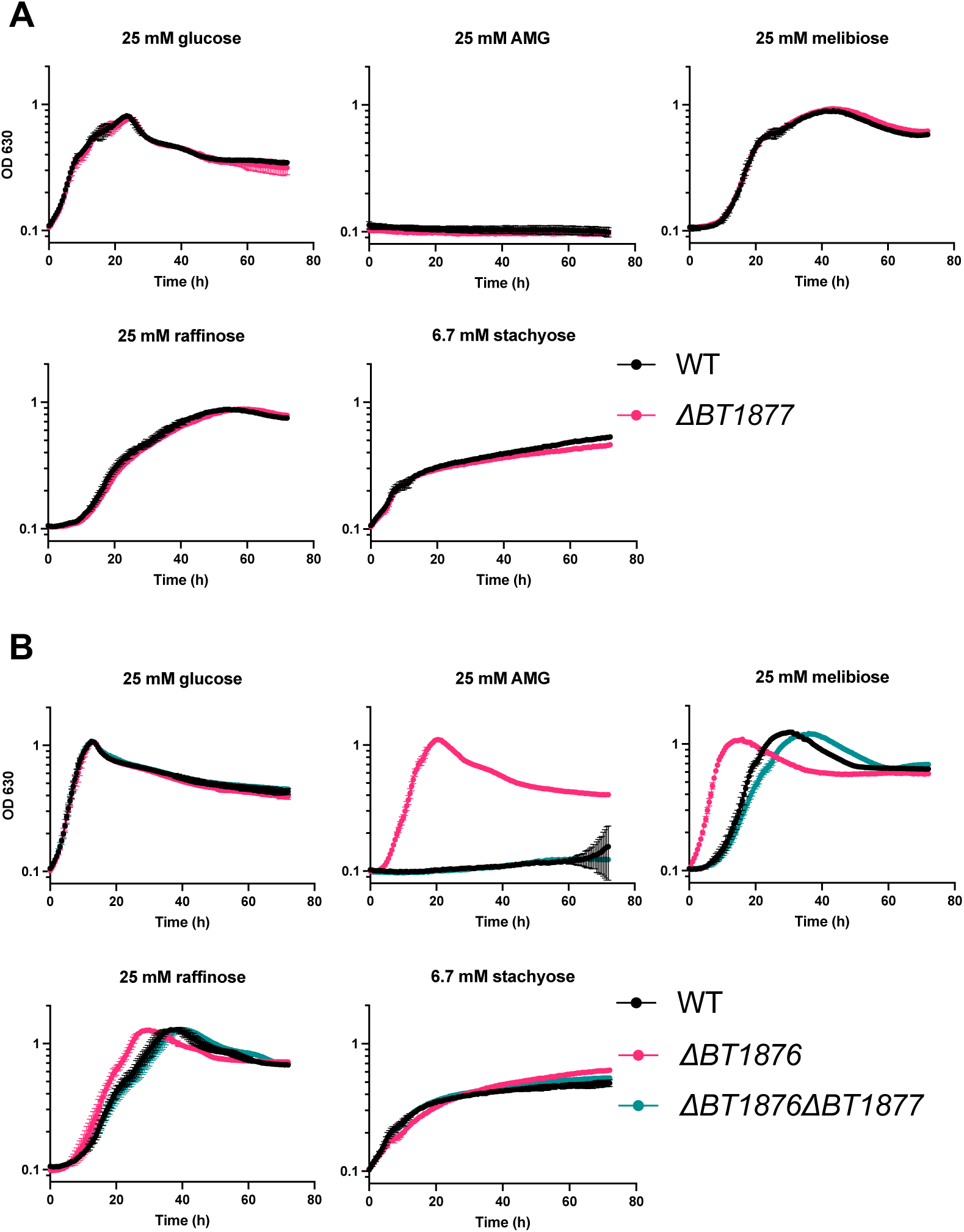
The PUL24 sigma factor *BT1877* is required for better growth of *BT1876* mutants on RFOs. (A) Growth curves of WT and *BT1877* mutant strain on RFOs. (B) Growth curves of WT, *BT1876* and *BT1876 BT1877* mutant strains on RFOs. In all panels, points and error bars represent the mean and SD of n=3 biological replicates. For each growth curve, the sugar used as the sole carbon source is indicated at the top along with the concentration used.

**Figure S7.**
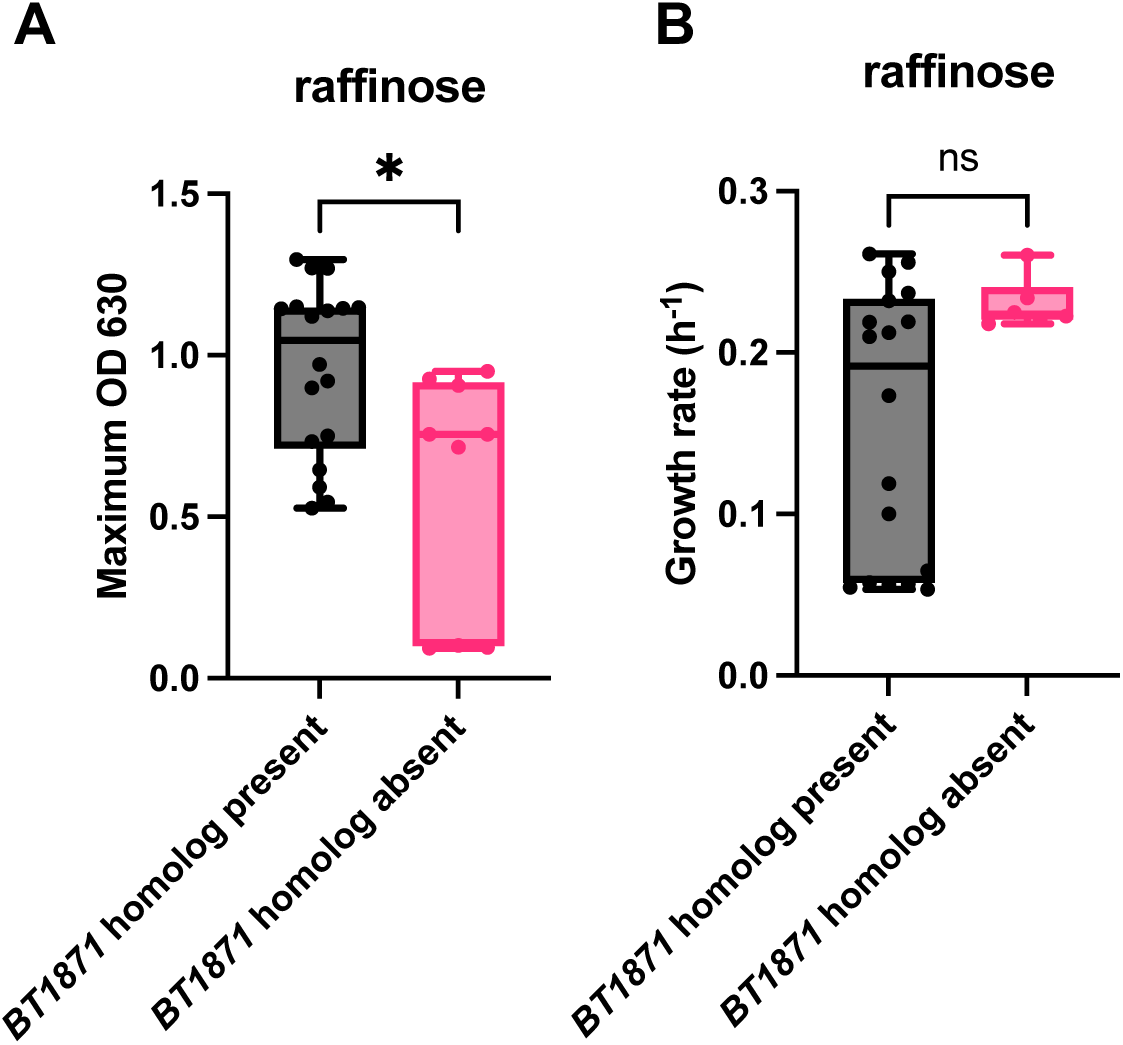
Differences in maximum ODs and growth rates on raffinose for *Bacteroides* species with or without a *BT1871* homolog. Boxplots showing comparison of maximum OD630 values (A) and growth rate (B) of *Bacteroides* species either with or without a *BT1871* homologue. Individual values were taken from growth curves depicted in 6A and 6B. Comparisons were done using a Mann Whitney U test and was found to be significant *, P<0.05 for (A) but not for (B).

**Figure S8.**
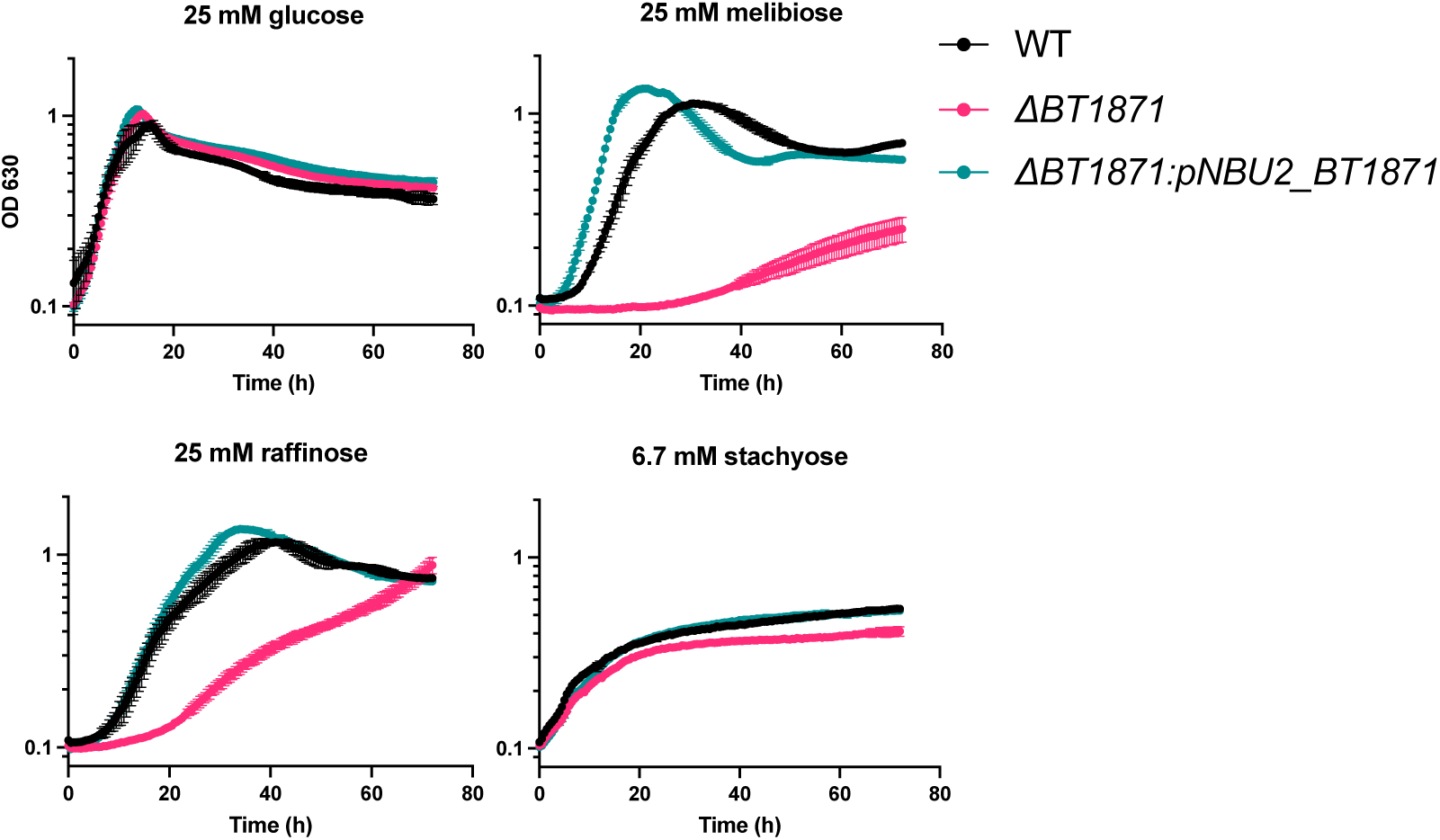
The constitutive *BT1871* expression cassette is active and restores growth on RFOs in a *BT1871* mutant strain. Growth curves of WT, *BT1871* mutant, and a *BT1871* mutant strain constitutively expressing *BT1871* under the control of the housekeeping sigma RpoD promoter (Δ*BT1871*:pNBU2_*BT1871*) on RFOs. Points and error bars represent the mean and SD of n=3 biological replicates. For each growth curve, the sugar used as the sole carbon source is indicated at the top along with the concentration used.

